# Karyotype stability and unbiased fractionation in the paleo-allotetraploid *Cucurbita* genomes

**DOI:** 10.1101/150755

**Authors:** Honghe Sun, Shan Wu, Guoyu Zhang, Chen Jiao, Shaogui Guo, Yi Ren, Jie Zhang, Haiying Zhang, Guoyi Gong, Zhangcai Jia, Fan Zhang, Jiaxing Tian, William J. Lucas, Jeff J. Doyle, Haizhen Li, Zhangjun Fei, Yong Xu

## Abstract

The *Cucurbita* genus contains several economically important species in the Cucurbitaceae family. Interspecific hybrids between *C. maxima* and *C. moschata* are widely used as rootstocks for other cucurbit crops. We report high-quality genome sequences of *C. maxima* and *C. moschata* and provide evidence supporting an allotetraploidization event in *Cucurbita*. We are able to partition the genome into two homoeologous subgenomes based on different genetic distances to melon, cucumber and watermelon in the Benincaseae tribe. We estimate that the two diploid progenitors successively diverged from Benincaseae around 31 and 26 million years ago (Mya), and the allotetraploidization happened earlier than 3 Mya, when *C. maxima* and *C. moschata* diverged. The subgenomes have largely maintained the chromosome structures of their diploid progenitors. Such long-term karyotype stability after polyploidization is uncommon in plant polyploids. The two subgenomes have retained similar numbers of genes, and neither subgenome is globally dominant in gene expression. Allele-specific expression analysis in the *C. maxima* × *C. moschata* interspecific F_1_ hybrid and the two parents indicates the predominance of *trans-*regulatory effects underlying expression divergence of the parents, and detects transgressive gene expression changes in the hybrid correlated with heterosis in important agronomic traits. Our study provides insights into plant genome evolution and valuable resources for genetic improvement of cucurbit crops.

## Background

Pumpkin, squash and gourd species (*Cucurbita* spp.,) belong to the Cucurbitaceae (cucurbit) family and consist of at least five domesticated and more than ten wild species^1,2^. *C*. *maxima* Duch. and *C*. *moschata* Duch. are two of the three most economically important cultivated *Cucurbita* species^3^. Archaeological data indicate that *C. maxima* and *C*. *moschata* originated in southern and northern South America, respectively, and were introduced to the Old World after European contact with the Americas^2^. *C. maxima* and *C*. *moschata* then underwent a great diversification in their China-Japan and India-Myanmar secondary domestication centers, respectively^2,4^. These *Cucurbita* crops are used as a staple food in many developing countries, and are consumed all over the world, mainly for their mature fruits and seeds, which are a rich source of nutritional compounds^2^. In addition to their culinary uses, the fruits are also used as ornaments and carved into decorative lanterns around Halloween. Nowadays, *C*. *maxima* and *C*. *moschata* are cultivated worldwide. The annual production of all *Cucurbita* species reached 25.2 million tons in 2014, with China and India leading in world production (http://faostat3.fao.org/).

The economic value of *C*. *maxima* and *C*. *moschata* is increasing, as they are also used as rootstocks for other cucurbit crops, including watermelon (*Citrullus lanatus*), cucumber (*Cucumis sativus*) and melon (*Cucumis melo*), to enhance tolerance to soilborne diseases and abiotic stresses^5^. The interspecific hybrid developed from a cross between *C. maxima* cv. Rimu and *C. moschata* cv. Rifu, ‘Shintosa’, is a popular rootstock for different cucurbits, and especially preferred in watermelon grafting for its Fusarium wilt resistance, cold-tolerance, and the ability to increase fruit weight, fruit quality and plant vigor^6^. *C. maxima*, in general, has better fruit flavor and texture than *C. moschata*, but is not as resistant as *C. moschata* to diseases and abiotic stresses. Interest in interspecific crossing between *C. maxima* and *C. moschata* has centered on transferring high-quality flesh traits from *C. maxima* to *C. moschata* and on introducing resistance traits from *C. moschata*^7^. However, the molecular mechanisms underlying these agronomical traits in these *Cucurbita* interspecific hybrids remain largely unexplored.

Compared with other cucurbit genera, *Cucurbita* has a higher chromosome number^1^ (2n = 40). Early cytogenetic and isozyme studies as well as a recent synteny analysis suggest a possible whole-genome duplication in this genus^8-10^. However, genomic resources are very limited in *Cucurbita* and little information is available regarding their genome and gene evolution. Here, we present the genome sequences of *C*. *maxima* and *C*. *moschata*, which provide evidence for an ancient allotetraploidization event in these two genomes, likely involving a hybridization between two highly diverged diploid progenitors. In the same nucleus, the two distinct subgenomes have largely maintained the chromosomal structures of the progenitors. Both subgenomes have experienced similar levels of gene loss, and neither is globally dominant in gene expression levels, with many homoeologous genes having divergent expression patterns. The draft genome sequences of *C. maxima* and *C. moschata* enabled us to analyze the allele-specific expression in the interspecific F_1_ hybrid, ‘Shintosa’, and determine the regulatory mechanisms underlying the expression divergence between *C. maxima* and *C. moschata*, as well as gene expression changes occurring upon interspecific hybridization that are correlated with heterosis in the *Cucurbita* F_1_ hybrid.

## Results

### Genome assembly, anchoring and quality evaluation

The genomes of *C. maxima* cv. Rimu and *C. moschata* cv. Rifu, the two parents of the ‘Shintosa’ rootstock, were sequenced and assembled. A total of 109.3 and 80.1 Gb of high-quality cleaned Illumina paired-end and mate-pair reads were generated for *C. maxima* and *C. moschata*, respectively, representing 283× and 215× coverage of their genomes (**Table S1**). Based on the 17-mer depth distribution analyses of the sequenced reads (**Figure. S1**), the genome sizes of *C. maxima* and *C. moschata* were estimated to be 386.8 and 372.0 Mb, respectively. *De novo* assemblies resulted in a draft genome of 271.4 Mb for *C. maxima* and 269.9 Mb for *C. moschata* (Fig. 1; Figure. S2), representing 70.2% and 72.6% of their estimated genome sizes, respectively. The assembled *C. maxima* genome had scaffold and contig N50 sizes of 3.7 Mb and 40.7 kb, respectively, and the N50 sizes for the scaffolds and contigs of the *C. moschata* assembly were 4.0 Mb and 40.5 kb, respectively (**Table 1**).

**Figure 1.**
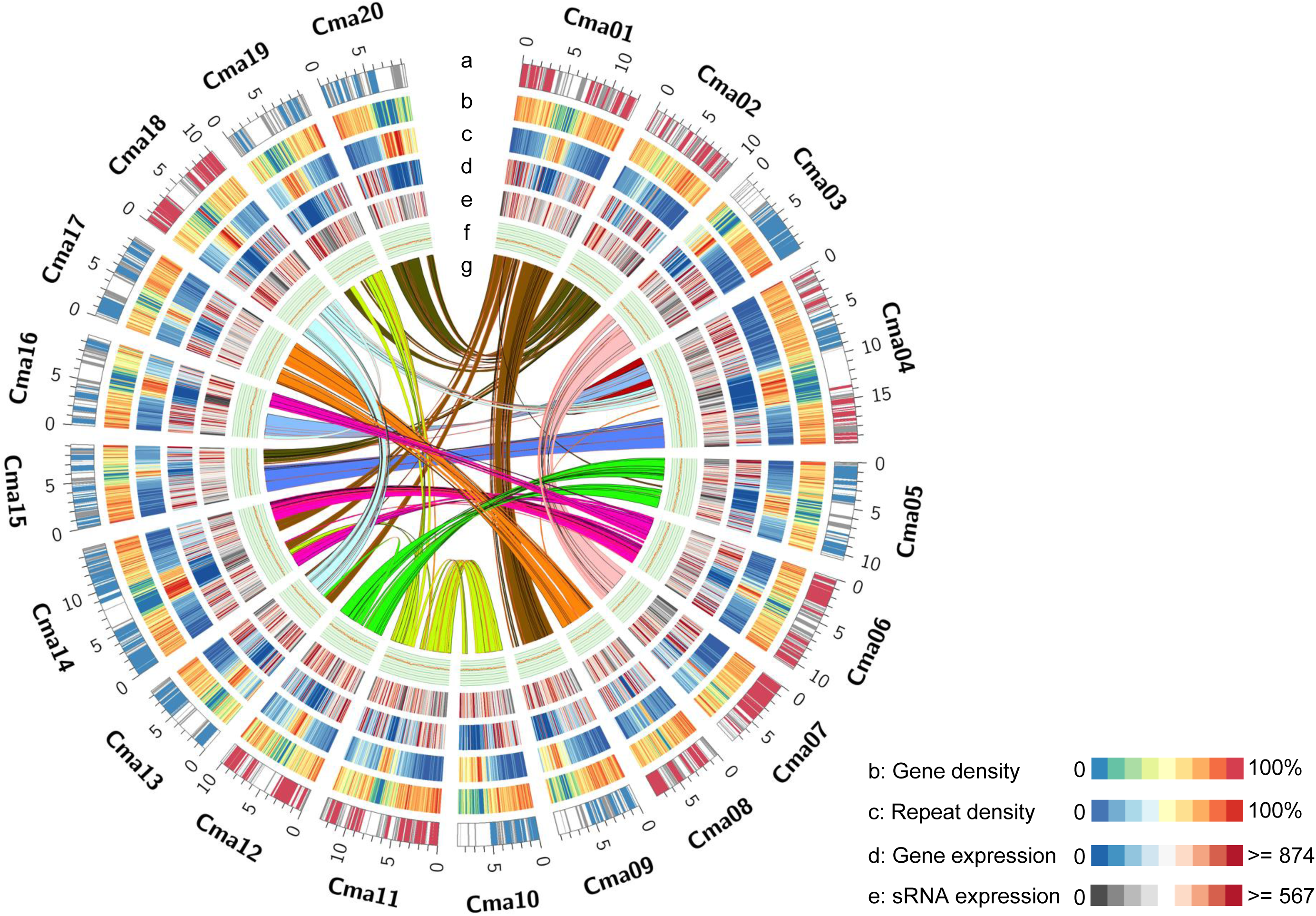
Genomic landscape of *C. maxima*. (**a**) Ideogram of the 20 *C. maxima* pseudochromosomes (in Mb scale). Syntenic blocks assigned to subgenomes A and B, and those that could not be assigned are labeled in red, blue and grey, respectively. (**b**) Gene density represented by percentage of genomic regions covered by genes in 200-kb windows. (**c**) Repeat density represented by percentage of genomic regions covered by repeat sequences in 200-kb windows. (**d**) Gene expression. Gene expression levels were estimated by fragment counts per million mapped fragments in 200-kb windows. (**e**) sRNA expression. sRNA expression levels were estimated by read counts per million mapped reads in 200 kb windows. (**f**) GC content in 200-kb windows. (**g**) Syntenic blocks depicted by connected lines.

**Table 1.** Summary of *C. maxima* and *C. moschata* assembled genomes

|  |  | Scaffold |  | Contig |  |
| --- | --- | --- | --- | --- | --- |
|  |  | Size (bp) | Number | Size (bp) | Number |
| *C. maxima* | Longest | 11,871,986 | 1 | 400,020 | 1 |
|  | N50 | 3,717,157 | 24 | 40,681 | 1,813 |
|  | Total | 271,413,401 | 8,299 | 265,448,559 | 25,524 |
| *C. moschata* | Longest | 11,258,782 | 1 | 292,205 | 1 |
|  | N50 | 3,995,720 | 24 | 40,480 | 1,897 |
|  | Total | 269,943,085 | 3,500 | 262,991,909 | 17,340 |

Contigs and scaffolds $\geq 500$ bp in length were included in the genome assemblies.

We constructed a *C. maxima* intraspecific genetic map containing 2,030 SNPs spanning 3,008.2 cM across 20 linkage groups (LGs) and a more saturated interspecific map (13,779 SNPs, 4,053.9 cM, 20 LGs) (**Table S2**). Both the intraspecific and interspecific maps and a previous map constructed using the same intraspecific population^11^ were largely collinear with all 20 linkage groups having shared scaffolds. Using the genetic maps of *C. maxima* (both the intra- and interspecific), 92 scaffolds (211.4 Mb, 77.9% of the assembly) were anchored to the 20 LGs, among which 64 scaffolds (201.2 Mb, 74.14 of the assembly) were oriented (**Figure. S3a**; **Table S3**). Likewise, we also constructed a *C. moschata* intraspecific map that contained 3,487 SNPs grouped into 20 LGs with a total genetic distance of 3,376.1 cM, and an interspecific map that had 16,214 SNPs distributed in 20 LGs and an overall genetic distance of 2,233.5 cM (**Table S2**). Based on these two genetic maps, 98 *C. moschata* scaffolds (238.5 Mb, 88.4% of the assembly) were anchored, and 71 scaffolds (221.3 Mb, 82.0%) were oriented (Figure. S3b; **Table S3**).

To evaluate the quality of our assemblies, we first aligned genome reads back to the assembled genomes. About 93% and 87% of the reads used in *de novo* assemblies were mapped back to the *C. maxima* and *C. moschata* genomes, respectively, in a proper paired-end relationship (**Table S4**). We then mapped RNA-Seq reads to the assembly, and of all RNA-Seq reads derived from different tissues of *C. maxima* (including fruit, leaf, stem and root), 93.8- 95.8% were mapped to the genome. Similarly, 86.2-95.0% of *C. moschata* RNA-Seq reads were mapped (**Table S5**). Further evaluation using BUSCO^12^ revealed that 97.0% and 97.3% of the core eukaryotic genes were covered by genomes of *C. maxima* and *C. moschata,* respectively, and 95.5% and 95.9% were completely covered (**Table S6**). In summary, the high coverage of the genome and RNA-Seq reads as well as the core eukaryotic genes indicated the high quality of the assembled *C. maxima* and *C. moschata* genomes.

### Repeat sequence prediction and gene annotation

Repeat sequences accounted for more than 40% of both *C. maxima* and *C. moschata* genome assemblies (**Table S7**). Among these repeats, 69.9% in *C. maxima* and 62.9% in *C. moschata* were annotated as long terminal repeat (LTR) retrotransposons, and predominantly, the *copia*- and *gypsy*-type LTRs. Similar findings have been reported in the genomes of other species in the Cucurbitaceae, including cucumber^13^, melon^14^ and watermelon^15^.

We predicted 32,076 and 32,205 protein-coding genes in the *C. maxima* and *C. moschata* genomes, respectively, much higher than numbers of genes predicted in genomes of cucumber 16 (23,248), melon^14^ (27,427) and watermelon^15^ (23,440). Biological functions were assigned to 26,993 (84.2%) genes in *C. maxima* and 27,230 (84.6%) in *C. moschata*. A total of 2,414 and 2,336 transcription factors were identified in *C. maxima* and *C. moschata*, respectively, numbers that are close to double those in the other sequenced cucurbit genomes (**Table S8**). We identified 30 and 57 disease resistance genes encoding nucleotide-binding site leucine-rich repeat (NBS-LRR) proteins in *C. maxima* and *C. moschata*, respectively (**Table S9**). Low copy number of disease resistance genes is known for other cucurbit genomes^17^. Orthology and synteny analyses between *C. maxima* and *C. moschata* showed that most *C. maxima* NBS-LRR genes had at least one orthologue in *C. moschata*, while certain genes were unique to *C. moschata* (**Table S9**).

### Two distinguishable paleo-subgenomes with nearly intact progenitor chromosome structures in *C. maxima* and *C. moschata*

We identified 195 paralogous syntenic blocks (7,773 homoeologous gene pairs) in *C. maxima* and 170 blocks (7,742 gene pairs) in *C. moschata*, with synonymous substitution rate (*K*s) values peaking at 0.32 (**Fig. 2a**; **Figure. S4** and **S5**), collectively covering 92.5% and 94.1% of the gene space, respectively. These homoeologous pairs clearly indicated that both *C. maxima* and *C. moschata* genomes underwent a whole-genome duplication event that was not observed in other sequenced cucurbits, including cucumber^13^, melon^14^, watermelon^15^ and bitter gourd (*Momordica charantia*) (**Fig. 2a**).

**Figure 2.**
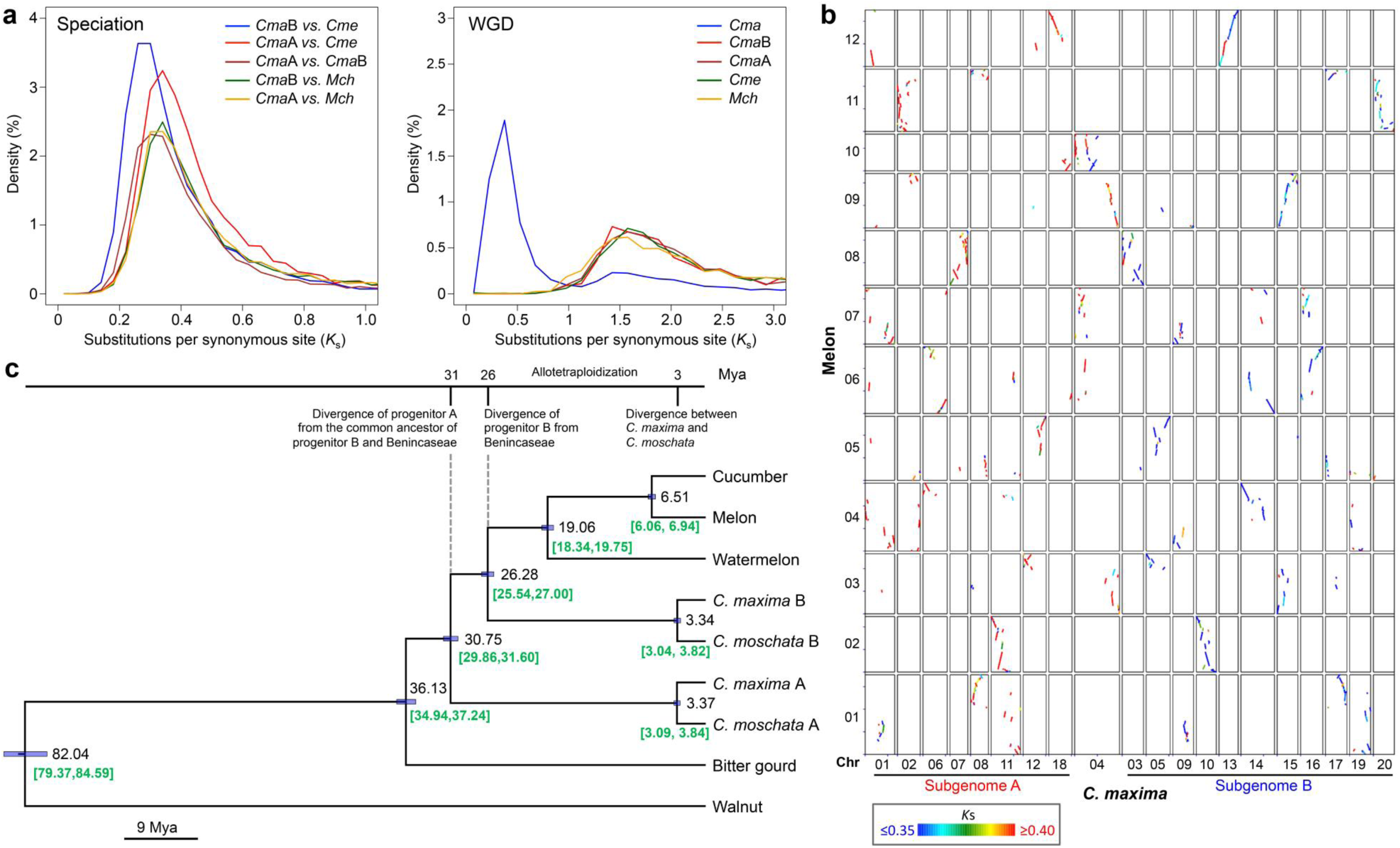
Genome evolution of *Cucurbita*. (**a**) Distribution of *K*s between orthologous or paralogous genes in the genomes of *C. maxima* (*Cma*; subgenomes A and B, *Cma*A and *Cma*B), melon (*Cucumis melo*, *Cme*) and bitter gourd (*Momordica charantia*, *Mch*). WGD, whole genome duplication. (**b**) Comparison between the genomes of *C. maxima* and melon. Two syntenic regions in *C. maxima* corresponding to one melon region with different *C. maxima*-watermelon *K*s values are marked by different colors. (**c**) Phylogenetic tree and timing of the speciation events. Estimated divergence times are shown at the nodes. The 95% highest probability density intervals are indicated as node bars and numbers in green listed below.

Whole-genome synteny analysis revealed pairs of *C. maxima* (or *C. moschata*) homoeologous regions shared between chromosomes corresponding to two subgenomes (**Fig. 1**; **Figure. S4**). Unlike the three subgenomes of *Brassica rapa*, which cannot be differentiated based on the *Arabidopsis thaliana*-*B. rapa K*s values^18^, the two subgenomes of *Cucurbita* could be distinguished according to their different genetic distances to their orthologues in the Benincaseae species, including melon, cucumber and watermelon (**Fig. 2b**; **Figure. S6** and **S7**). About 63.2% 58.7% and 68.3% of the *C. maxima* genomic regions (64.0%, 62.2% and 69.5% for *C. moschata*) syntenic to melon, cucumber and watermelon, respectively, were composed of homoeologous blocks with significantly different *Cucurbita*-Benincaseae *K*s values. Except for chromosome 4, all other chromosomes contained only homoeologous blocks with either higher or lower *K*s values (Fig. 2b; **Table S10**).

Based on these findings, we separated the entire chromosomes into two groups to represent the two paleo-subgenomes. Eight and 11 chromosomes were assigned to subgenomes A and B, respectively. Chromosome 4 was divided into three segments with two assigned to subgenome A and one to subgenome B (**Figure. S8**). The reconstructed subgenomes A and B of *C. maxima* contained 14,816 and 16,068 protein-coding genes, respectively (15,136 and 16,473 genes in *C. moschata*, respectively). Phylogenetic analysis clearly indicated an allotetraploid origin of *Cucurbita* from two progenitors that diverged from one another approximately 30.75 million years ago (Mya) (**Fig. 2c**). One ancestor gave rise to watermelon, cucumber and melon in the Benincaseae tribe, as well as to the B-genome progenitor of *Cucurbita*, which has not been sampled and may be extinct. No extant diploid species that could represent the A-genome progenitor have been sampled, so it is possible that this genome donor is also extinct. The oldest possible date for the polyploidy event is the date of divergence of Benincaseae from the B-genome of *Cucurbita* (approximately 26.28 Mya, **Fig. 2c**). There was extensive shared synteny between the two *Cucurbita* species with only a few inversions (**Figure. S9**), consistent with a shared polyploidy event. The divergence date of the two *Cucurbita* species (3.04-3.84 Mya, **Fig. 2c**) provides a minimum age for the allopolyploidy event. There was no significant difference of *K*s values between *C. maxima*-*C. moschata* subgenome A and subgenome B orthologous gene pairs (**Figure. S10**), suggesting that the two subgenomes evolved with similar mutation rates after polyploidization. Furthermore, the distribution of *Cucurbita*-Benincaseae syntenic blocks with higher (or lower) *Ks* along an entire chromosome (except for chromosome 4) (**Fig. 2b**) indicated a low level of large-scale inter-chromosomal exchanges since the allotetraploidization event, which suggested karyotype stability in the polyploid *Cucurbita* genomes. This was further strongly supported by the fact that the same ancestral states of four chromosomes from progenitor B (*Cucurbita* chromosomes 3, 10, 13 and 15) were also found in the modern melon genome (**Fig. 2b**).

### Genome and gene evolution after allotetraploidization

Following polyploidization, functionally redundant duplicated (homoeologous) genes often revert to single-copies through fixation of null mutations and fractionation mediated by intrachromosomal recombination, unless prevented by mechanisms such as gene dosage-balance constraints and functional diversification^19^. Recent studies have shown that gene loss after polyploidization can be biased towards one of the progenitor genomes, such as in Arabidopsis^20^, maize^21^, *Brassiaca rapa*^18^ and cotton^22^. We analyzed patterns of gene loss in *C. maxima* and *C. moschata* after the genome merger using high-confidence retained homoeologous genes and singletons (whose duplicated copies were lost) that had syntenic orthologues in both cucumber and watermelon. Unlike in the aforementioned plant species, in *Cucurbita* we observed random loss of genes (unbiased fractionation) from the two subgenomes. Seven pairs of putative homoeologous chromosomes retained similar numbers of genes, whereas slightly biased gene-loss patterns were detected for Chr08-Chr17 and Chr18-Chr04, 13, 16 pairs (**Fig. 3a**; **Figure. S11**; **Table S11**).

**Figure 3.**
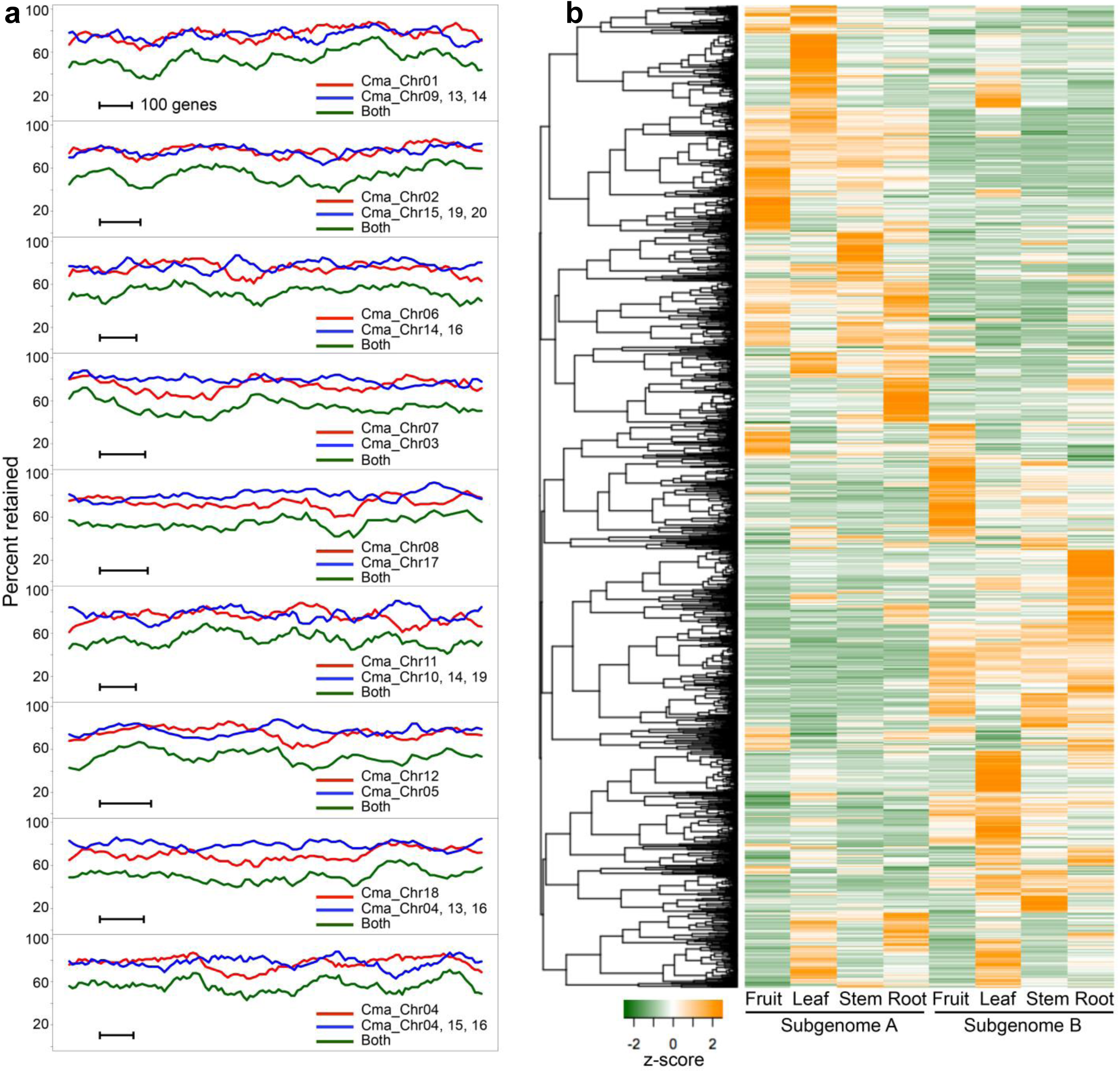
Gene retention and functional diversification of duplicated genes after allotetraploidization. (**a**) Fractionation in pairs of homoeologous regions in *C. maxima*. Average gene retention percentages are shown in a 100-gene window along each subgenome. Retention of genes in subgenomes A, B and both are shown in red, blue and green, respectively. (**b**) Expression profiles of high-confidence homoeologous gene pairs in the two subgenomes of *C. maxima*. FPKM values of each pair were standardized to have a mean of zero and standard deviation of one across all the samples for the clustering analysis.

A total of 6,550 genes in *C. maxima* (6,639 in *C. moschata*) were assigned to 348 KEGG pathways, as expected, most of which contained similar numbers of genes from the two subgenomes. However, we did find that pathways of pentose and glucuronate interconversions, galactose metabolism, and starch and sucrose metabolism were composed of more subgenome B genes, and phenylalanine metabolism had more subgenome A genes (**Table S12**). The biased numbers were due to more genes from subgenome B encoding four enzymes, polygalacturonase (K01184), *β*-glucosidase (K01188), *β*-fructofuranosidase (K01193) and galacturan 1,4- *α* - galacturonidase (K01213) shared by the three carbohydrate metabolism pathways, and more genes from subgenome A encoding caffeoyl-CoA O-methyltransferase (K00588), tyrosine aminotransferase (K00815) and phenylalanine ammonia-lyase (K10775) in phenylalanine metabolism (**Table S12;** **Figure S12**).

We next tested the presence or absence of genome dominance by comparing expression levels between pairs of high-confidence duplicated genes. Using RNA-Seq data from fruit, leaf, stem and root, we found no evidence of overall biased expression between the two subgenomes (**Figure. S13**). This result should be expected given the lack of biased fractionation, and given that bias in gene expression is thought to be responsible for biased homoeologous gene loss^23^.

Among all the high-confidence homoeologous genes in *C. maxima*, 11,296 (5,648 pairs) were retained in two copies and 4,063 were singletons. Similarly, the *C. moschata* genome retained 11,572 duplicated genes (5,786 pairs) that resulted from the allotetraploidization, and 5,027 had become single-copy genes. Most of these retained duplicates (4,507 pairs) and singletons (3,633 genes) were shared between the two species, which is to be expected considering the relatively recent divergence of *C. maxima* and *C. moschata*. In addition to almost double the number of transcription factors in *Cucurbita* compared to watermelon, cucumber and melon (**Table S8**), enriched gene ontology (GO) categories for retained duplicates included response to unfolded protein, G-protein coupled receptor signaling pathway, and steroid hormone mediated signaling pathway (**Table S13**). The retention of these genes could be due to dosage-balance constraints since such genes often function in macromolecular complexes or regulatory cascades, and are prone to be co-retained to avoid detrimental consequences of disrupting stoichiometric relationships among interacting partners^19,24,25^. On the other hand, genes functioning in DNA and RNA metabolic processes were over-represented in the singletons (**Table S13**), consistent with previous findings on under-retention of genes with conserved biological functions^24,25^.

Evolutionary innovations in allopolyploids are provided by combining parental legacies and novel divergence in regulatory elements and/or coding regions and rewiring of duplicated regulatory networks upon, or subsequent to polyploidization^26,27^. For most duplicated genes, the specific expression pattern of a gene was not shown in its homoeologue (Fig. 3b; Figure. S14). About 49.0% of high-confidence duplicated gene pairs in *C. maxima* (48.6% in *C. moschata*) were differentially expressed by at least 2-fold. These results suggest extensive variation in gene expression patterns between the two subgenomes, which might well be caused by *cis*-regulatory divergence between the diploid progenitors, immediate alteration upon genome merger, as well as long-term evolutionary forces^28,29^.

Robust evidence of positive selection was detected between 49 pairs of duplicated genes in *C. maxima* and also 49 pairs in *C. moschata* (**Table S14**), which might have acquired new functions important for adaptation and/or increased *Cucurbita* fruit morphological complexity. Among these genes, 23 pairs were common between *C. maxima* and *C. moschata*, suggesting functional diversification of some homoeologous genes before the speciation of *C. maxima* and *C. moschata*. These genes included four in *C. maxima* and *C. moschata* encoding OVATE family proteins, whose expansion and functional divergence are present in many plant species, and members of this family are key regulators of fruit shape in tomato and pepper^30^, consistent with the highly diverse fruit shapes of *Cucurbita* species.

After polyploidization, retained duplicated genes may exhibit increased expression divergence between species compared to the single-copy genes^31^. We compared gene expression levels between *C. maxima* and *C. moschata*, and found that the proportions of differentially expressed genes were slightly higher for duplicated genes (**Table S15**). We detected positive selection between 1,784 pairs of orthologous genes in *C. maxima* and *C. moschata* (**Table S16**), among which 948 had high-confidence retained duplicates in both genomes (out of 9,109 pairs of such genes in the two genomes, 10.4%) and 301 pairs (out of 3,622, 7.8%) were singletons. These results suggested the duplicates might have contributed more to the functional divergence of orthologous genes between *C. maxima* and *C. moschata*.

### *Cucurbita* lineage-specific gene expansion

Using protein sequences of *C. maxima*, *C. moschata*, three Benincaseae species (cucumber, melon and watermelon), grape and papaya, we detected 52 gene families that underwent significant expansion in the *Cucurbita* lineage (**Table S17**). These included genes encoding pectin lyases (OG0000048, OG0000060 and OG0000491), pathogenesis-related 1 proteins (OG0000041), terpene synthases (OG0000056), carbohydrate esterases (OG0001297), pectinesterase inhibitors (OG0000281), subtilisin-like serine protease inhibitors (OG0000148) and mitogen-activated protein kinases (OG0001303) that are predicted to function in plant cell wall modification during plant growth and fruit softening, and/or plant defense. The expanded defense-related genes could provide alternative defense mechanisms complementary to the low numbers of NBS-LRR genes in *C. maxima* and *C. moschata*. We also detected expansions of gene families that may control meristem activity and organ morphogenesis such as orthologues of *Arabidopsis LONELY GUY* genes (OG0000097) and *CLAVATA3/ESR-RELATED 45* (OG0002365), *RADIALIS-LIKE SANT*/*MYB* genes (OG0000105), *ROTUNDIFOLIA*-*like* genes (OG0000137) and genes encoding NAC domain-containing proteins (OG0010138), CONSTANS-like proteins (OG0003099) and Gibberellin 3 beta-hydroxylase family proteins (OG0011381). The higher retention rate of these genes after small- and/or large-scale duplications suggested that they might underlie important adaptive traits during evolution to develop survival strategies. While most expanded families contained genes from both subgenomes, the expansions of some families were contributed by only one subgenome (Figure. S15; **Table S17**). For example, the pectin lyase (OG0000048) and terpenene synthase (OG0000056) genes were almost all from subgenome B, and the pectinesterase (OG0000281) and carbonhydrate esterase (OG0001297) genes were nearly exclusively from subgenome A. This might result from gene loss and/or tandem duplication specific to one subgenome before and/or after genome merger.

### Expression divergence between *C. maxima* and *C. moschata*

Divergence in the regulatory mechanisms controlling gene expression is pivotal in evolution by providing the source of phenotypic diversity between divergent species. Successful interspecific hybridization between *C. maxima* and *C. moschata* offered an opportunity to evaluate the *cis*-and *trans*- contributions to their expression divergence through quantifying the total and allele-specific gene expression levels in the interspecific F_1_ hybrid ‘Shintosa’ and the two parents. About 22-32% of the genes (7,303 in fruit, 6,464 in leaf, 7,749 in stem and 9,371 in root) were differentially expressed between *C. maxima* and *C. moschata* (**Table S18**). Of these genes, 62% to 80% (4,528 in fruit, 4,691 in leaf, 6,223 in stem and 7,210 in root) were found to have only *trans*-regulatory effects. The remaining genes were classified as having only *cis*-effects (24.0%, 19.3%, 13.2% and 13.9% of the differentially expressed genes in fruit, leaf, stem and root, respectively) or both *cis*- and *trans*-effects (14.0%, 8.2%, 6.5% and 9.2%). This result suggested that *trans*-regulatory effect was the predominant cause of expression divergence between *C. maxima* and *C. moschata*. A general trend has been identified from previous studies that *trans*-effects account for more of the expression differences between less divergent species, whereas *cis*-effects primarily cause expression variation between more genetically distinct species^32-34^. The prevalence of *trans*-variation between *C. maxima* and *C. moschata* is consistent with genetic similarity due to their relatively recent divergence. Among the 1,784 pairs of *C. maxima*-*C. moschata* orthologous genes under significant positive selection (see above), 351 pairs were found to have significant *cis*-effects, indicating gene functional diversification accompanied by regulatory divergence.

### Gene expression alteration in the interspecific F_1_ hybrid

Interspecific crosses play an important role in *Cucurbita* breeding for transferring favorable traits between species^2^. Hybridization between *C. maxima* and *C. moschata* created novel interactions between the parental alleles and a new regulatory environment in the F_1_ hybrid. Genes exhibiting dominant and transgressive expression patterns might underlie heterotic phenotypes and combined favorable characters from each parent, such as high carotenoid levels in fruit from *C. maxima*^35^, and better insect pest resistance and abiotic stress tolerance from *C. moschata*^36,37^. In the F_1_ hybrid, 4,002 (fruit), 6,718 (leaf), 6,732 (stem) and 7,067 (root) genes were expressed higher than or equal to the high parent (up-regulated), or lower than or equal to the low parent (down-regulated), representing 12.5-22.0% of the total genes. Among these genes, 49.6-75.0% (2,417 in fruit, 3,330 in leaf, 5,049 in stem and 4,590 in root) exhibited *C. moschata*-like expression, and 7.7-19.0% showed *C. maxima*-like expression (**Fig. 4a, b**). About 41.7-50.8% of the genes exhibiting dominant and transgressive expression patterns had significant *trans*-regulatory effects in different tissues, whereas 6.3-20.1% had significant *cis*-effects (**Fig. 4c, d**).

**Figure 4.**
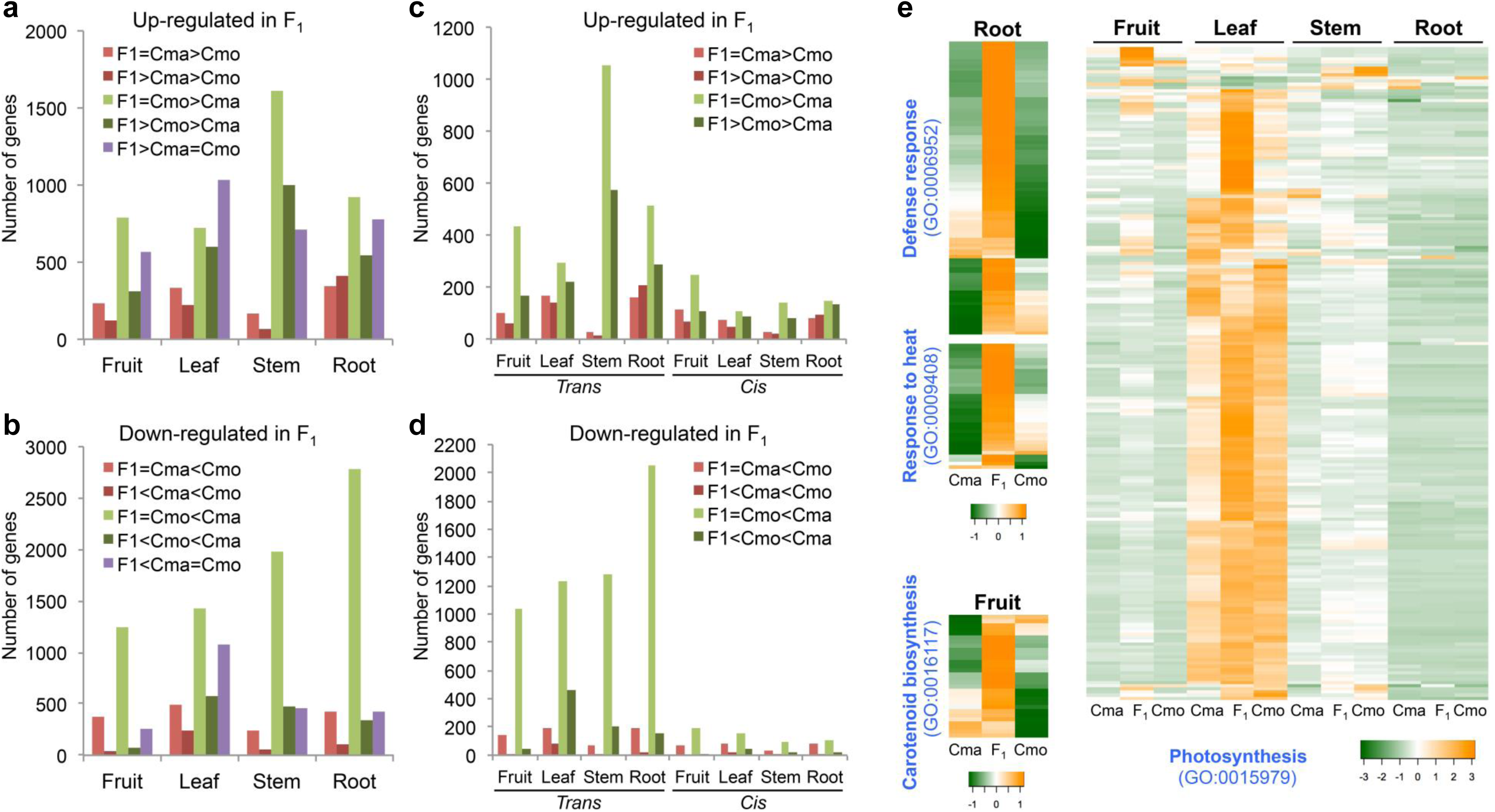
Regulatory mode of dominantly and transgressively expressed genes in the interspecific F_1_ hybrid, ‘Shintosa’. (**a**-**b**) Numbers of up- (**a**) or down-regulated (**b**) genes in F_1_ compared to the parents. (**c**-**d**) Numbers of up- (**c**) or down-regulated (**d**) genes with significant *trans*- or *cis*- regulatory effects. (**e**) Expression patterns of genes in selected over-represented biological processes. FPKM values of each gene were standardized to have a mean of zero and standard deviation of one across compared samples. Cma, *C. maxima*. Cmo, *C. moschata*.

We identified GO terms enriched in the dominantly and transgressively expressed genes in the hybrid. In the hybrid root, enrichment of genes involved in defense response (GO:0006952) and response to heat (GO:0009408) was found among up-regulated genes (**Fig. 4e**), which could be related to the elevated stress tolerance of ‘Shintosa’. Interestingly, down-regulated genes in the hybrid root were significantly enriched with pathways related to active root growth, including auxin-signaling (GO:0009734), auxin-transport (GO:0060918), cell division (GO:0051301), cell growth (GO:0016049), cell wall organization/biogenesis (GO:0071554), and root hair cell development (GO:0080147) (**Table S19**), suggesting a tradeoff between defense response and growth processes, which has also been observed in Arabidopsis^38,39^. We also found that carotenoid biosynthesis (GO:0016117) was significantly over-represented in the up-regulated genes in the fruit flesh of the hybrid. A comparison of gene expression in the carotenoid biosynthetic pathway among the two parents and F_1_ showed that the expression levels of phytoene synthase (PSY) and lycopene *β*-cyclase (LYCB) genes were higher in the F_1_ fruit compared to the parents (**Table S20;** Figure. S16). The duplicated copies of *PSY* (and *LYCB*) were homoeologues resulting from the allotetraploidization. Only one of the homoeologues was highly expressed in either *C. maxima* or *C. moschata* fruit, whereas both copies were highly expressed in the F_1_ hybrid (**Figure. S16**). The *β*-carotene hydroxylase (CHYB) genes were expressed at a relatively lower level in *C. moschata* fruit compared with the F_1_ hybrid (higher) and *C. maxima* (highest), which could lead to different levels of hydroxylation of *α*- and *β*-carotene. Another transgressively expressed carotenoid biosynthesis gene in the F_1_ fruit encoded zeaxanthin epoxidase (ZEP), which epoxidizes zeaxanthin to violaxanthin. The photosynthesis-related pathways (GO:0015979 and GO:0009765) were significantly over-represented in all three aerial tissues in the up-regulated genes (**Fig. 4e**). These photosynthesis-related genes included those encoding light-harvesting chlorophyll a/b binding proteins and subunits of photosystem I and II, previously shown to be associated with heterosis in other plant species^40^.

## Discussion

Most economically important genera in the Cucurbitaceae belong to the Cucurbiteae and Benincaseae tribes^41^. We present high-quality draft genome sequences of two *Cucurbita* species of the Cucurbiteae, *C. maxima* and *C. moschata*, as well as evidence for an ancient allotetraploidy event in *Cucurbita*. The divergence of the two diploid progenitors occurred around 31 Mya, soon after the common ancestor of Cucurbiteae and Benincaseae diverged from the Momodiceae that includes bitter gourd. Although the hybridization date of the two diploid progenitors is uncertain, the divergence between *C. maxima* and *C. moschata* is estimated to be at 3 Mya, constraining the tetraploidization to be earlier than this date.

Our results suggest only limited inter-chromosomal rearrangements since the hybridization of the two diploid progenitors. Chromosome structural variation between the two subgenomes is therefore likely due to different fission and fusion events in the two progenitor genomes before hybridization, with the current *Cucurbita* subgenomes largely maintaining the chromosome structures of the two progenitors despite sharing the same nucleus for at least 3 million years. Such karyotype stability is in contrast to the frequent homoeologous exchanges among subgenomes of allotetraploid cotton during 1-2 million years after polyploidization^42^ and considerable rearrangements experienced by many other plant polyploids^43^, but has been observed in amphibians such as allotetraploid frog (*Xenopus laevis*) that originated from an ancient hybridization event 17-18 Mya^44^, as well as in newly synthesized allotetraploid wheat with genome combinations analogous to natural tetraploids^45^. The karyotype stability in the *Cucurbita* genomes could be due to divergence between the two diploid genome donors, which prevented meiotic pairing of homoeologous chromosomes and subsequent exchanges. Karyotype stability could play a key role in the initial establishment of the *Cucurbita* species by avoiding fitness costs associated with genomic aberrations immediately following polyploidization.

The two subgenomes have retained similar numbers of genes, and neither subgenome is dominant in overall expression levels. Fractionation bias and genome dominance have been observed in several plant species after polyploidization, including Arabidopsis, maize, rice, *Brassica rapa* and diploid cotton^46^. On the other hand, lack of genome dominance and biased fractionation in the genomes of banana, poplar and soybean was reported by Garsmeur et al.^47^, who hypothesized that this different postpolyploidy subgenome evolution pattern was possibly associated with ancient autopolyploidy events, contradicting the hypothesis that soybean is an allotetraploid^48^. Here, in *Cucurbita*, we identified another case of random fractionation and lack of genome-wide expression bias after an ancient allopolyploidy event.

The genome sequences of *C*. *maxima* and *C*. *moschata* allowed us to perform an allele-specific gene expression analysis in the interspecific F_1_ hybrid, to parse the genome-wide expression differences between *C. maxima* and *C. moschata* into *cis-* and *trans-* regulatory effects, and detect transgressive gene expression changes in the hybrid correlated with increased disease resistance and growth vigor. The expression patterns of carotenoid biosynthesis genes (including *PHY*, *LYCB*, *CHYB* and *ZEP*) in the fruits of the F_1_ hybrid and the parents suggest that the hybrid harnesses the active pathway components used by both parents. Our result is consistent with previous findings that the predominant carotenoids in *C. moschata* fruits are *α*- and *β*-carotene, whereas *C. maxima* accumulates lutein and violaxanthin at high levels^35,49^, and also suggests that interspecific hybridization between *C. maxima* and *C. moschata* could lead to a different fruit carotenoid profile and potentially higher total carotenoid content.

## Conclusions

High-quality genome sequences of two *Cucurbita* species, *C. maxima* and *C. moschata*, were assembled using high coverage of Illumina paired-end and mate-pair reads, and each constructed into 20 pseudochromosmes. In addition to providing evidence of a whole-genome duplication in the *Cucurbita* lineage, our comparative genomics analyses revealed the evolutionary history of this ancient allotetraploidy event, including successive divergence of the two diploid progenitors (probably now extinct) of *Cucurbita* from the common ancestor of *Cucumis* and *Citrullus* species that happened around 31 and 26 Mya, and hybridization of the diploids earlier than 3 Mya. The genomes of *Cucurbita* represent a type of paleo-allotetraploid genomes with stable karyotypes that maintained the chromosome structures of the diploid genome donors. We also showed that the *Cucurbita* genomes underwent unbiased fractionation after allotetraploidization associated with the lack of genome dominance. Allelic-specific expression analysis in the interspecific hybrid of *Cucurbita* and the two parents showed that the expression divergence between *C. maxima* and *C. moschata* was predominantly controlled by *trans*-regularoty effects. We also detected transgressive gene expression changes in the hybrid correlated with heterosis in important agronomic traits such as disease resistance and carotenoid biosynthesis. The genome sequences of *C. maxima* and *C. moschata* have improved our knowledge on the genome evolutionary of plant polyploids, and provide an invaluable resource for the study of agronomic traits in *Cucurbita* and genetic improvement of the species.

## Materials and methods

### Plant materials and DNA/RNA extraction

Two inbred *Cucurbita* lines, *C. maxima* cv. Rimu and *C. moschata* cv. Rifu, were selected for genome sequencing. Seedlings were grown in the greenhouse under natural light supplemented with artificial light (16/8 h light/dark) and transferred to a dark room for 24 hours to promote starch degradation. Young leaves were harvested and stored at −80°C. To construct genetic maps, two intraspecific F2 populations were generated from crosses between *C. maxima* cultivars, Rimu and SQ026, and between *C. moschata* cultivars, Rifu and Honey jujube. The interspecific population was derived from a *C. maxima* cv. Rimu × *C. moschata* cv. Rifu F_1_ individual backcrossing to *C. maxima* cv. Rimu. For transcriptome sequencing, fully developed leaves, stems, roots, and fruit flesh at 46 days after pollination were collected from *C. maxima* cv. Rimu, *C. moschata* cv. Rifu and their F_1_ hybrid ‘Shintosa’. Three independent biological replicates were prepared for each sample.

Genomic DNA and total RNA were extracted using the QIAGEN DNeasy Plant Mini Kit and the QIAGEN RNeasy Plant Mini Kit, respectively, following the manufacturer’s instructions (QIAGEN, Valencia, California, USA). DNA and RNA quality was evaluated via agarose gel electrophoresis and their quantity was determined on a NanoDrop (Thermo Fisher Scientific, Waltham, MA, USA).

### Genomic and RNA-Seq library construction and sequencing

Paired-end libraries with insert sizes ranging from 200 bp to 1 kb and mate-pair libraries with insert size ranging from 3 to 15 kb were prepared using Illumina Genomic DNA Sample Preparation kit and the Nextera Mate Pair Sample Preparation kit (Illumina, San Diego, CA), respectively, following the manufacturer’s instructions. Strand-specific RNA-Seq libraries were prepared from total RNA following the protocol described by Zhong et al.^50^. Genomic and RNA-Seq libraries were sequenced on the Illumina HiSeq 2500 system with the paired-end mode.

### *De novo* genome assembly

Raw Illlumina reads were processed to collapse duplicated read pairs into unique pairs. Duplicated read pairs were defined as those having identical bases at positions of 14 to 50 in both left and right reads. The non-duplicated reads were further processed with Trimmomatic^51^ to remove adaptors and low-quality sequences. Read pairs from mate-pair libraries were processed with the ShortRead package^52^ to remove junction adaptors, followed by removing reads shorter than 40 bp and error correction with QuorUM^53^.

The high-quality cleaned reads were assembled into scaffolds with SOAPdenovo2^54^ and gaps in the resulting scaffolds were filled with the GapCloser program in the SOAPdenovo2 package. Pilon^55^ was used to improve the assembly by correcting bases, fixing mis-assemblies and further filling gaps. Potential contaminations from microorganisms were detected by aligning the assemblies to NCBI non-redundant nucleotide (nt) database using BLASTN with an e-value cutoff of 1e-5. Scaffolds with more than 90% of their length similar to bacterial sequences were considered contaminants and removed. Finally, scaffolds contained within other scaffolds with sequence identity >99% and coverage >99% were removed.

### Genetic map construction and scaffold anchoring

To anchor the assembled *C. maxima* and *C. moschata* scaffolds, we constructed high-density genetic maps using two intraspecific F2 populations (186 *C. maxima* and 186 *C. moschata* individuals) and one interspecific F_1_BC_1_ population (186 individuals) derived from *C. maxima* Rimu × SQ026, *C. moschata* Rifu × Honey jujube, and *C. maxima* Rimu × *C. moschata* Rifu, respectively. For the interspecific F_1_BC_1_ population, *C. maxima* Rimu was used as the recurrent parent. Genomic DNA was extracted from young healthy leaf tissues of F_2_ and F_1_BC_1_ individuals and the parents using the cetyltrimethylammonium bromide (CTAB) method. DNA concentration was measured using an ND-2000 spectrophotometer (NanoDrop, Wilmington, DE, USA) and quality was assessed by electrophoresis using 1% agarose gels with a lambda DNA standard. Genotyping of these plants was performed following the GBS protocol^56^, using *ApeKI* as the restriction enzyme. The resulting libraries were multiplexed and sequenced on a HiSeq 2500 system (Illumina Inc. USA) with single-end mode and read length of 101 bp. The GBS sequencing reads were processed using the TASSEL-GBS (v4) pipeline^57^ for SNP calling. Briefly, for each population, the reads from all samples were combined and collapsed into a master tag list. Tags that occurred at least 10 times were retained and mapped to the *C. maxima* or *C. moschata* genome using BWA^58^ (v0.7.12) with default parameters. Alignments with mapping quality ≥ 1 were used for SNP calling. The minimum genotype quality was set to 10. SNPs with significant segregation distortion (Chi-square test, *p* <0.05), high data missing rate (>30%) or low minor allele frequency (MAF <0.03) were removed. The resulting SNPs were used to construct genetic maps using the minimum spanning tree (MST) algorithm^59^ implemented in the R package ASMap (https://cran.r-project.org/web/packages/ASMap/). During scaffold anchoring, one initial *C. moschata* scaffold was identified as chimeric, on the basis of the same scaffold mapping to different linkage groups and the lack of mate-pair read support at the point of potential misjoining. This scaffold was split into two new ones.

### Annotation of transposable elements

A *de novo* long terminal repeat retrotransposon (LTR-RT) library and a miniature inverted repeat transposable elements (MITE) library were constructed by screening the assembled *C. maxima* and *C. moschata* genomes using LTRharvest^60^ and MITE-Hunter^61^, respectively. Both genomes were then masked using RepeatMasker (http://www.repeatmasker.org/) with the LTR-RT and MITE libraries. The unmasked sequences in the genomes were further searched for repeat elements using RepeatModeler (http://www.repeatmasker.org/RepeatModeler.html). All the repetitive sequences generated above were combined into a single repeat library for each genome, and then compared against the Swiss-Prot database^62^. Sequences that matched non-TE proteins in the database were removed from the repeat libraries. The remaining TEs were classified using REPCLASS^63^. The classified repeat libraries were then used to identify TEs in the assembled *C. maxima* and *C. moschata* genomes with RepeatMasker (http://www.repeatmasker.org/).

### Protein coding gene prediction and functional annotation

The repeat-masked *C. maxima* and *C. moschata* genomes were used for gene prediction with MAKER^64^, which combines evidences from *ab initio* gene prediction, transcript mapping and protein homology to define the confident gene models. SNAP^65^ and AUGUSTUS^66^ were used for *ab initio* gene predictions. RNA-Seq data were assembled using Trinity ^67^ with the *de novo* mode and the genome-guided mode, respectively. The two assemblies were combined and aligned to the assembled genomes by the PASA2 pipeline^68^. The resulting alignments were used as the transcript evidence. Protein sequences from Arabidopsis^69^, watermelon^15^, cucumber^16^, and melon^14^ and the UniProt (Swiss-Prot plant division) database were aligned to the genomes using Spaln^70^ to provide the protein homology evidence. The same annotation pipeline was also applied to the bitter gourd genome^71^, which predicted 20,778 protein-coding genes that were used in the analyses in this study.

For gene annotation, protein sequences of the predicted genes were compared against the Arabidopsis protein and UniProt (Swiss-Prot and TrEMBL) databases using BLAST, as well as the InterPro database using InterProScan^72^. Gene ontology (GO) annotations were obtained using Blast2GO^73^. Transcription factors were identified using the iTAK program^74^.

### Detection of lineage-specific expansion of protein families

Lineage-specific gene family expansion was inferred from orthologous groups using the software CAFE^75^ (v3.0). Orthologous groups were constructed using OrthoFinder^76^ with protein sequences from grape, papaya, *C. maxima*, *C. moschata* and three other cucurbit species (watermelon, cucumber and melon). The random gene birth and death rates were estimated across the species tree composed of the seven species using the maximum likelihood method. Orthologous groups with accelerated rate of expansion in the *Cucurbita* genus (represented by *C. maxima* and *C. moschata*) were determined with a threshold *p*-value of 0.05.

### Detection of positive selection

Positive selection on pairs of paralogous (within *C. maxima* or *C. moschata*) or orthologous genes (between *C. maxima* and *C. moschata*) was detected using the CodeML application in the PAML package^77^. Positive selection was estimated by pairwise calculation of nonsynonymous/synonymous rate ratio (*K*a/*K*s or *ω*) between all members of an orthologous group using the codon-based ‘branch’ model^78^. Substitution rates were averaged across a gene (option ‘NSsites = 0’). The tested paralogous or orthologous branch was used as the ‘foreground’ and all other cucurbit branches (cucumber, watermelon and melon) as the ‘background’. We first tested whether the *ω* of the paralogous (or orthologous) genes was different from the rest of the tree by comparing a two-ratio model (option ‘model=2’ allowing *ω* to vary among branches) to the null model 0 (assuming an identical ω among all branches). To further test whether the difference was due to positive selection or just relaxation of selection pressure, the two-ratio model result was contrasted against a model of neutrality (option ‘fix_omega =1’). If the model allowing different *ω* (in the first comparison) or positive selection (in the second comparison) fit the data significantly better, as judged by the likelihood-ratio test (LRT), difference in *ω* or positive selection was inferred. Twice the difference in log-likelihood values between two models was compared to a χ^2^ distribution with one-degree of freedom to determine the *p*-value.

### Analyses of genome evolution, phylogeny and divergence time

To identify syntenic regions within and between genomes, protein sequences of *C. maxima*, *C. moschata*, watermelon, cucumber, melon and bitter gourd were aligned against themselves or each other using BLASTP, and high-confidence collinear blocks were determined using MCScanX^79^ with an E-value cutoff of 1e-10. *K*s values of homologous pairs were calculated using the Yang-Nielsen algorithm implemented in the PAML package^80^. To construct the species tree containing the diploid progenitors of *C. maxima* and *C. moschata* (represented by the two subgenomes), watermelon, cucumber and walnut, we used 187 orthologous groups containing only one syntenic gene from each species. The CDS sequences of these genes had at least 80% coverage to their walnut orthologues, and were longer than 300 bp. The alignments of the 187 orthologous groups were used to construct phylogenetic trees with PhyML^81^ (v3.0; default parameters). A consensus tree was produced using the TreeAnnotator program in BEAST2^82^ with default parameters. The divergence time was estimated using the program BPP, under the multispecies coalescent model in a Bayesian framework with the fixed species phylogeny constructed above^83^. Based on the fossil information on the divergence time between Fagales and Cucurbitales, which is around 84 Mya^84^, the gamma prior G(2, 8.66) were assigned to the population size parameters (θs), with mean 2/8.66 = 0.2309, and the gamma prior G(282, 2445) was assigned to the divergence time at the root of the species tree (τ0). The analysis was run twice to confirm consistency between runs.

### RNA-Seq analysis

Raw RNA-Seq reads were processed by Trimmomatic^51^ (version 0.32) to remove adapter, polyA/T tails and low quality (quality score < 20) sequences. Reads equal to or longer than 40 bp were kept and aligned to the SILVA rRNA database (release 111) (https://www.arb-silva.de/) to remove rRNA contaminations. Cleaned reads were aligned to the *C. maxima* and *C. moschata* genomes using HISAT2^85^ allowing no mismatch. Alignment results were used to classify each read as *C. maxima*-specific, *C. moschata*-specific, or common to both species. Raw counts for each gene were then derived and normalized to fragments per kilobase of exon model per million mapped reads (FPKM). DESeq^86^ was used to identify differentially expressed genes between *C. maxima* and *C. moschata* orthologues with a cutoff of adjusted *p*-value < 0.005, fold-change >2 and FPKM ≥ 3 in at least one sample.

### *cis* and *trans* effects and inheritance mode classification

The orthologous relationship between a pair of *C. maxima* and *C. moschata* genes was detected with LAST^87^ by looking for unique best corresponding regions between the genomes of *C. maxima* and *C. moschata*. Gene pairs with more than 90% coverage in both genomes were selected for the allele-specific expression analyses. Statistical significance of *cis* and *trans* effects were determined using binomial and Fisher’s exact tests as described previously^88^. Briefly, DESeq^86^ analyses were carried out to identify genes that were expressed differently between the parents and/or had expression ratios of *C. maxima* and *C. moschata* alleles significantly different (adjusted *p*-value < 0.005) from 1 in the F_1_ hybrid. The identified genes were then subjected to a Fisher’s exact test to determine whether the expression ratios of *C. maxima* and *C. moschata* in the F_1_ were significantly different (adjusted *p*-value < 0.005) from the ratios between the parents. Based on the results from the binomial tests and Fisher’s exact tests, each gene was classified into one of the seven different regulatory categories, namely, *cis*, *trans*, *cis* + *trans*, *cis* × *trans*, compensatory, conserved and ambiguous, as described by McManus et al.^88^. The mode of inheritance for each gene was determined based on the magnitude of the expression level difference between the hybrid and parents. Genes with expression level in the hybrid deviated more than 1.25-fold from that of either parent were further classified as having additive, dominant, under-dominant or over-dominant expression.

### Data access

The Whole Genome Shotgun projects of *C. maxima* and *C. moschata* have been deposited at DDBJ/ENA/GenBank under the accessions NEWN00000000 and NEWM00000000, respectively. The versions described in this paper are NEWN01000000 and NEWM01000000. Raw genome and transcriptome sequence reads have been deposited into the NCBI sequence read archive (SRA) under accessions SRP106545, SRP106546 and SRP106360. The genome sequences and the annotations of *C. maxima* and *C. moschata* are also available at the Cucurbit Genomics Database (http://cucurbitgenomics.org).

## ACKNOWLEDGMENTS

This research was supported by grants from the Beijing Scholar Program (BSP026), the Beijing Excellent Talents Program (2014000021223TD03), the Ministry of Agriculture of China (CARS-26), the Beijing Natural Science Foundation (6141001 and 6144023), the US National Science Foundation (IOS-1339287, IOS-1339128 and IOS-1539831), and USDA National Institute of Food and Agriculture Specialty Crop Research Initiative (2015-51181-24285).

## AUTHOR CONTRIBUTIONS

Z.F., Y.X., S.W., J.J.D. and W.J.L. designed and managed the project. G.Z., H.L., S.G., Y.R., J.Z. H.Z., G.G., Z.J., F.Z., J.T and C.J. prepared samples and performed biological experiments for DNA sequencing and genetic map construction. H.S., S.W., C.J., S.G. and Y.R. performed data analyses. S.W. and H.S. wrote the manuscript. Z.F., J.J.D., W.J.L. and Y.X. revised the manuscript.

## COMPETING FINANCIAL INTERESTS

The authors declare no competing financial interests.

## Figure legends

**Figure S1.**
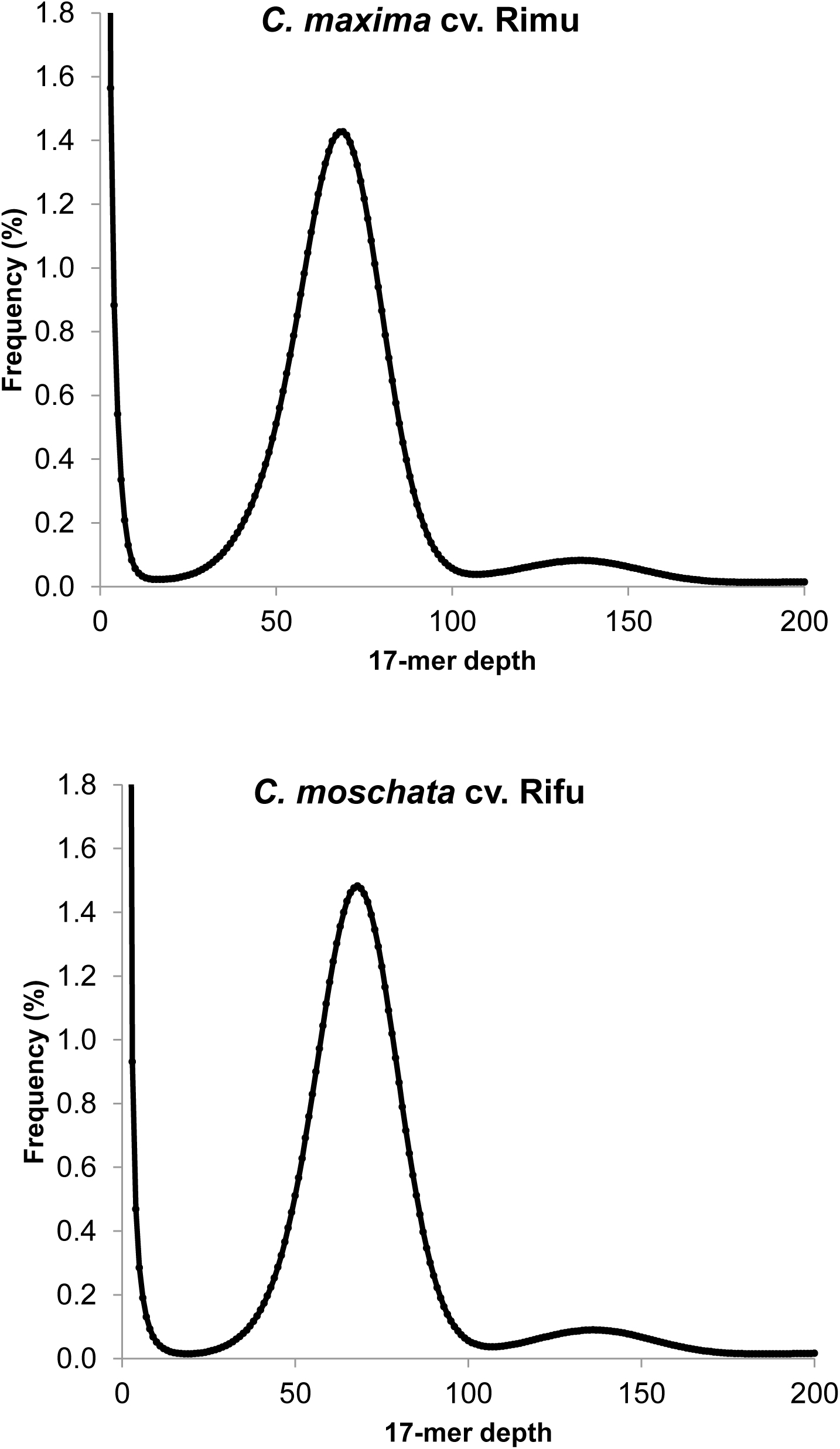
Kmer distribution of Illumina genomic sequencing reads of *C. maxima* and *C. moschata*.

**Figure S2.**
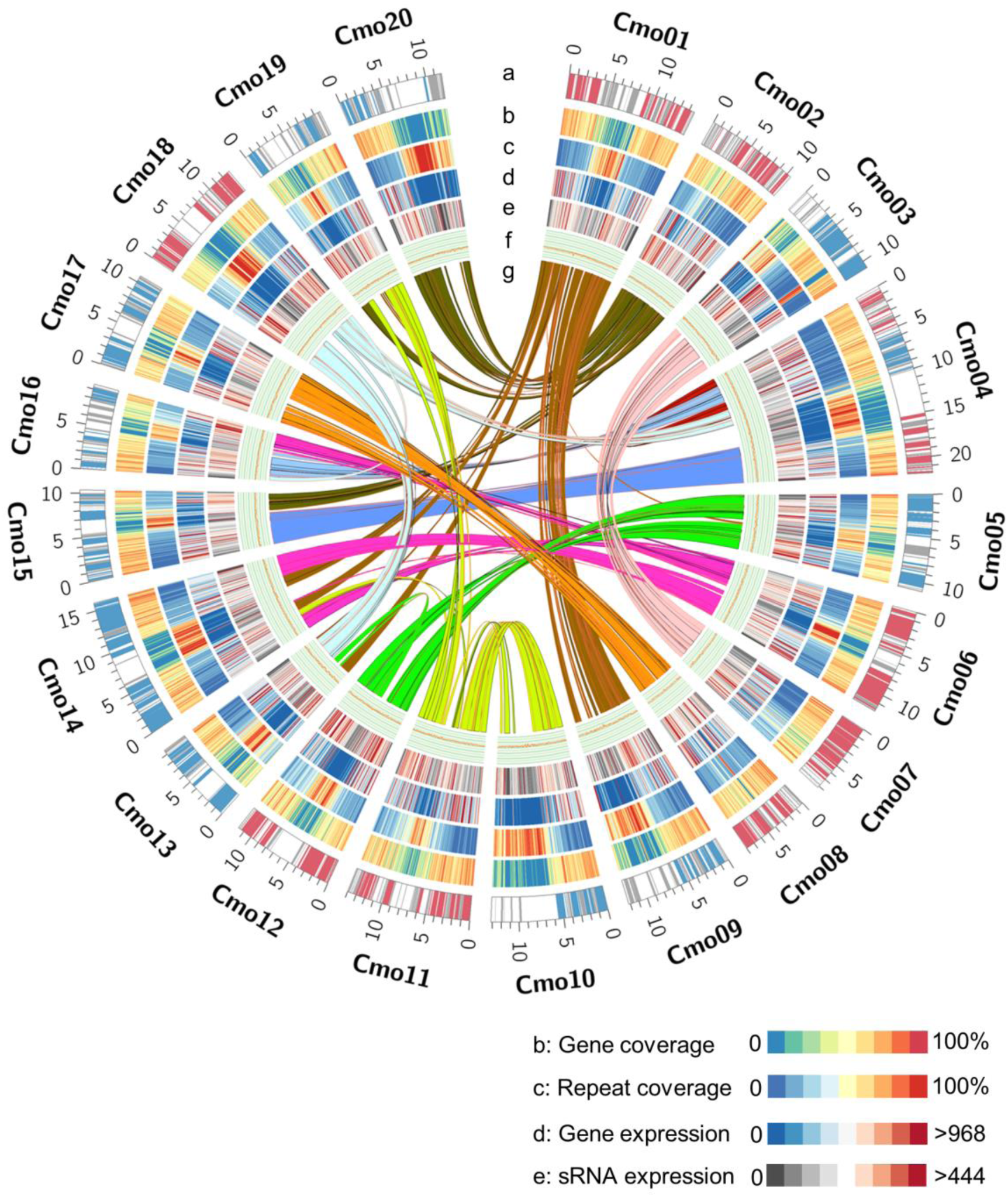
Genomic landscape of *C. moschata*. (**a**) Ideogram of the 20 *C. moschata* pseudochromosomes (in Mb scale). Syntenic blocks assigned to subgenomes A and B, and those that could not be assigned are labeled in red, blue and grey, respectively. (**b**) Gene density represented by percentage of genomic regions covered by genes in 200-kb windows. (**c**) Repeat density represented by percentage of genomic regions covered by repeat sequences in 200-kb windows. (**d**) Gene expression. Gene expression levels were estimated by fragment counts per million mapped fragments in 200-kb windows. (**e**) sRNA expression. sRNA expression levels were estimated by read counts per million mapped reads in 200-kb windows. (**f**) GC content in 200-kb windows. (**g**) Syntenic blocks depicted by connected lines.

**Figure S3.**
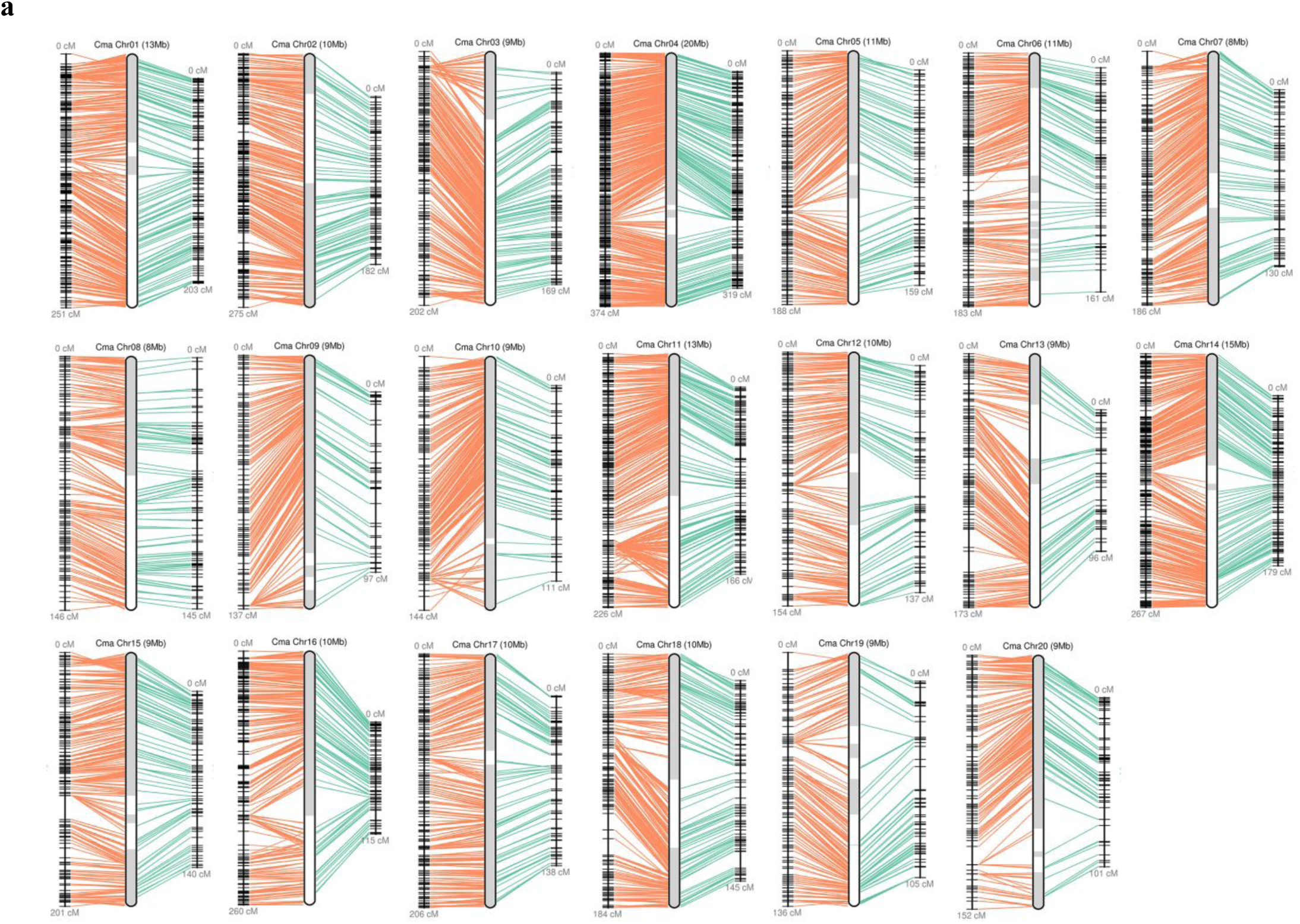
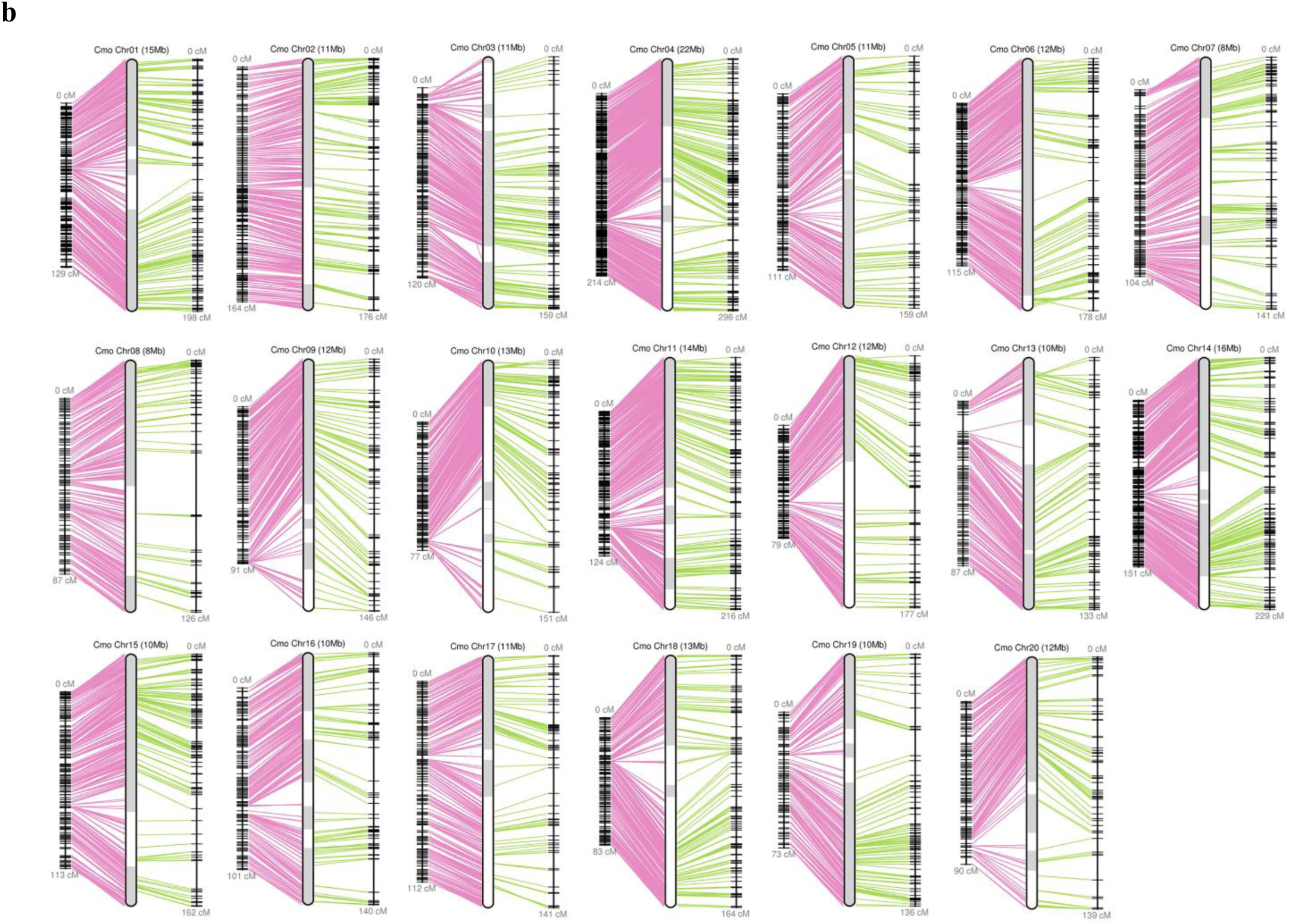
Anchoring *C. maxima* and *C. moschata* genome assemblies to high-density genetic maps. **(a)** *C. maxima* genome scaffolds (middle) were anchored to the linkage groups of the intraspecific *C. maxima* cv. Rimu × SQ026 (right) and interspecific *C. maxima* cv. Rimu × *C. moschata* cv. Rifu (left) populations. **(b)** *C. moschata* genome scaffolds (middle) were anchored to the linkage groups of the intraspecific *C. moschata* cv. Rifu × Honey jujube (right) and interspecific *C. maxima* cv. Rimu × *C. moschata* cv. Rifu (left) populations.

**Figure S4.**
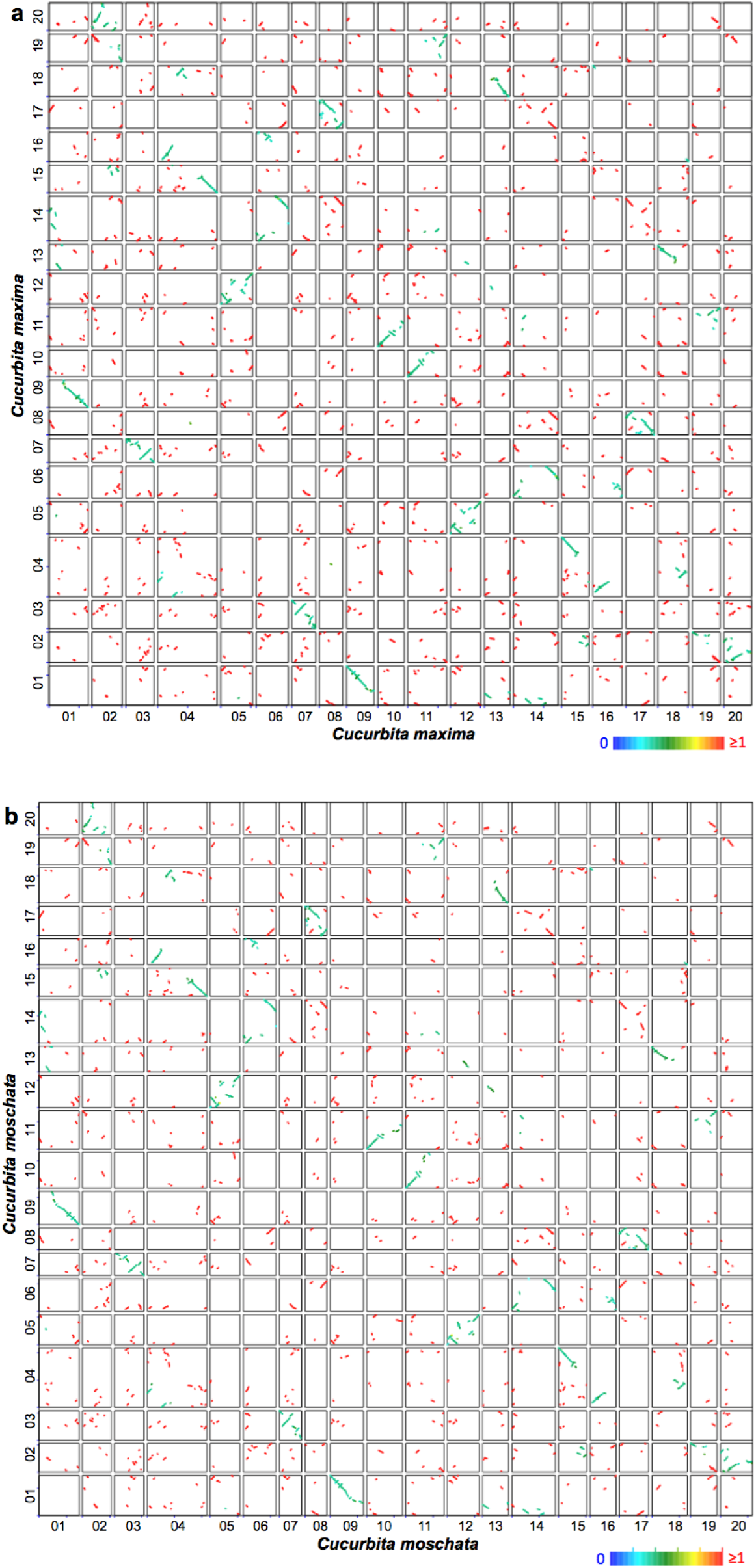
Syntenic dotplots showing the whole genome duplication in *C. maxima* (a) and *C. moschata* (b). Paralogous gene pairs are marked in colors based on the median *K*s values of syntenic blocks.

**Figure S5.**
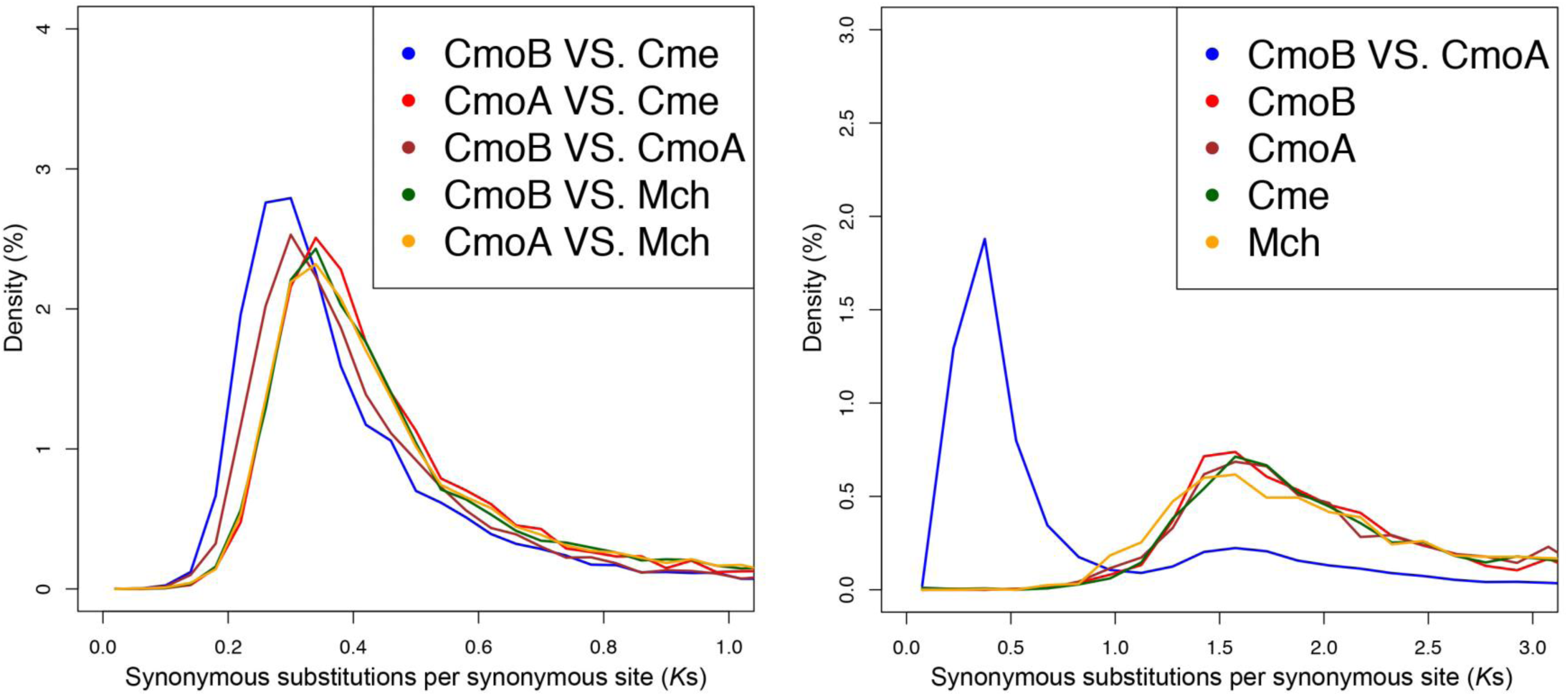
Distribution of *K*s between orthologous or paralogous genes in the genomes of *C. mochata* (Cmo; subgenome A and B, CmoA and CmoB), melon (*Cucumis melo*, Cme) and bitter gourd (*Momordica charantia*, Mch).

**Figure S6.**
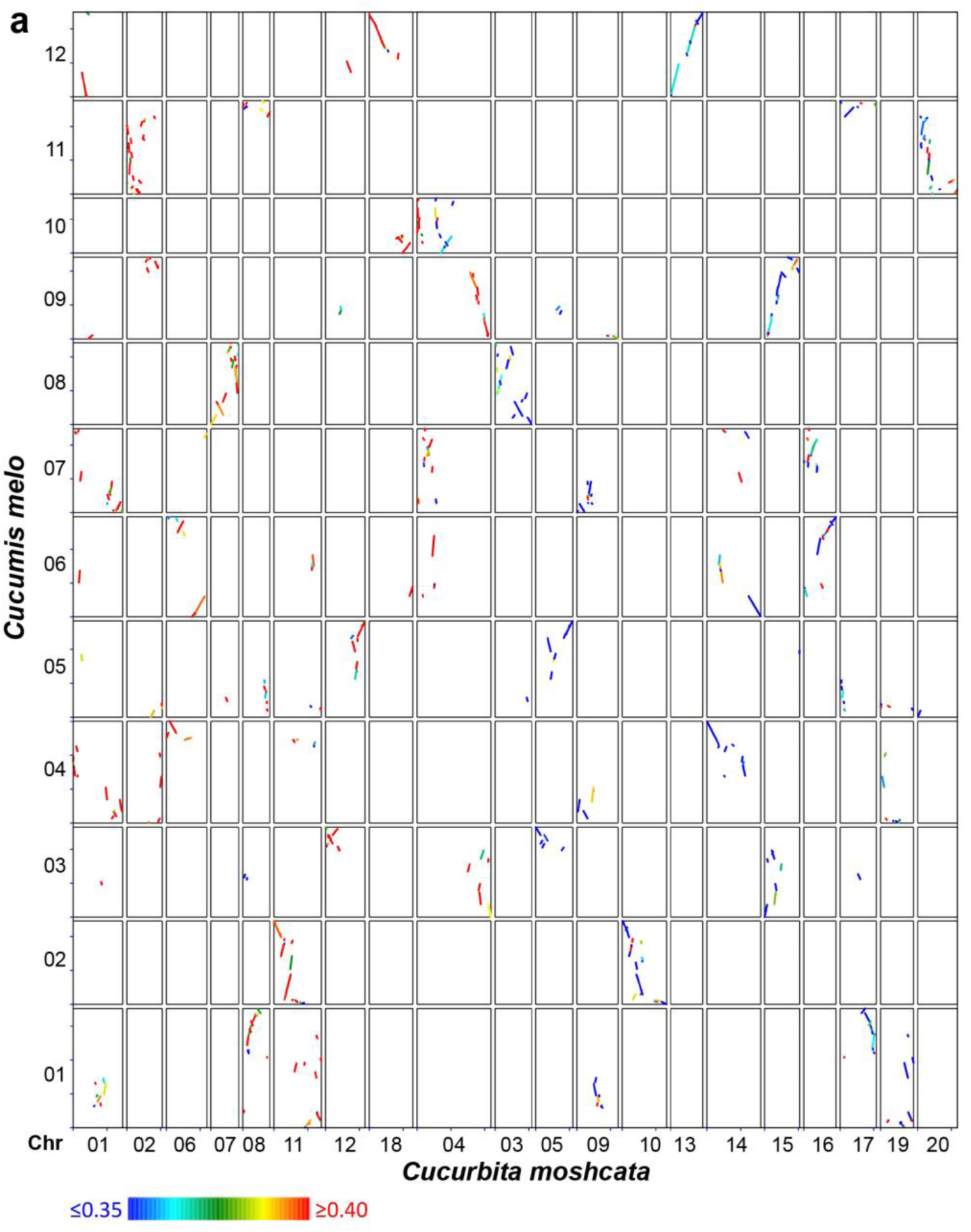
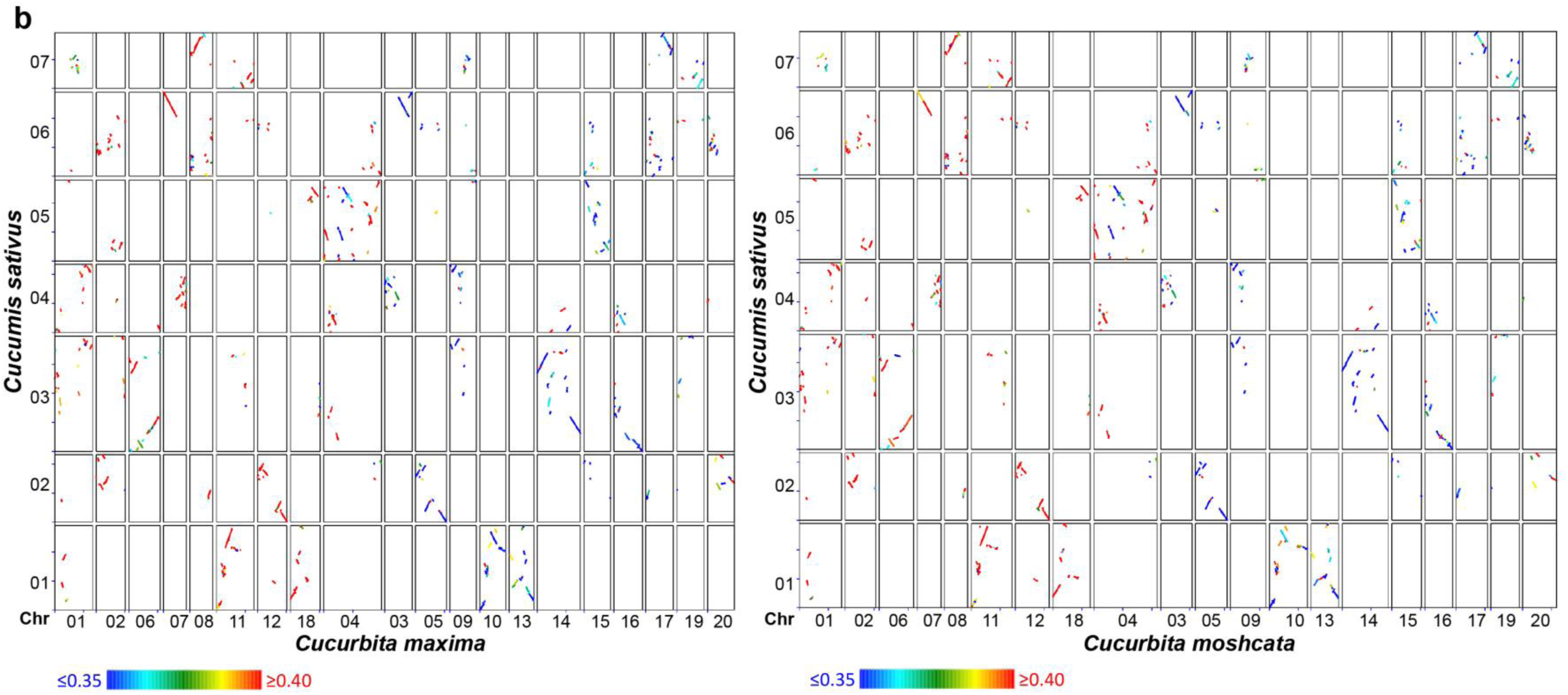
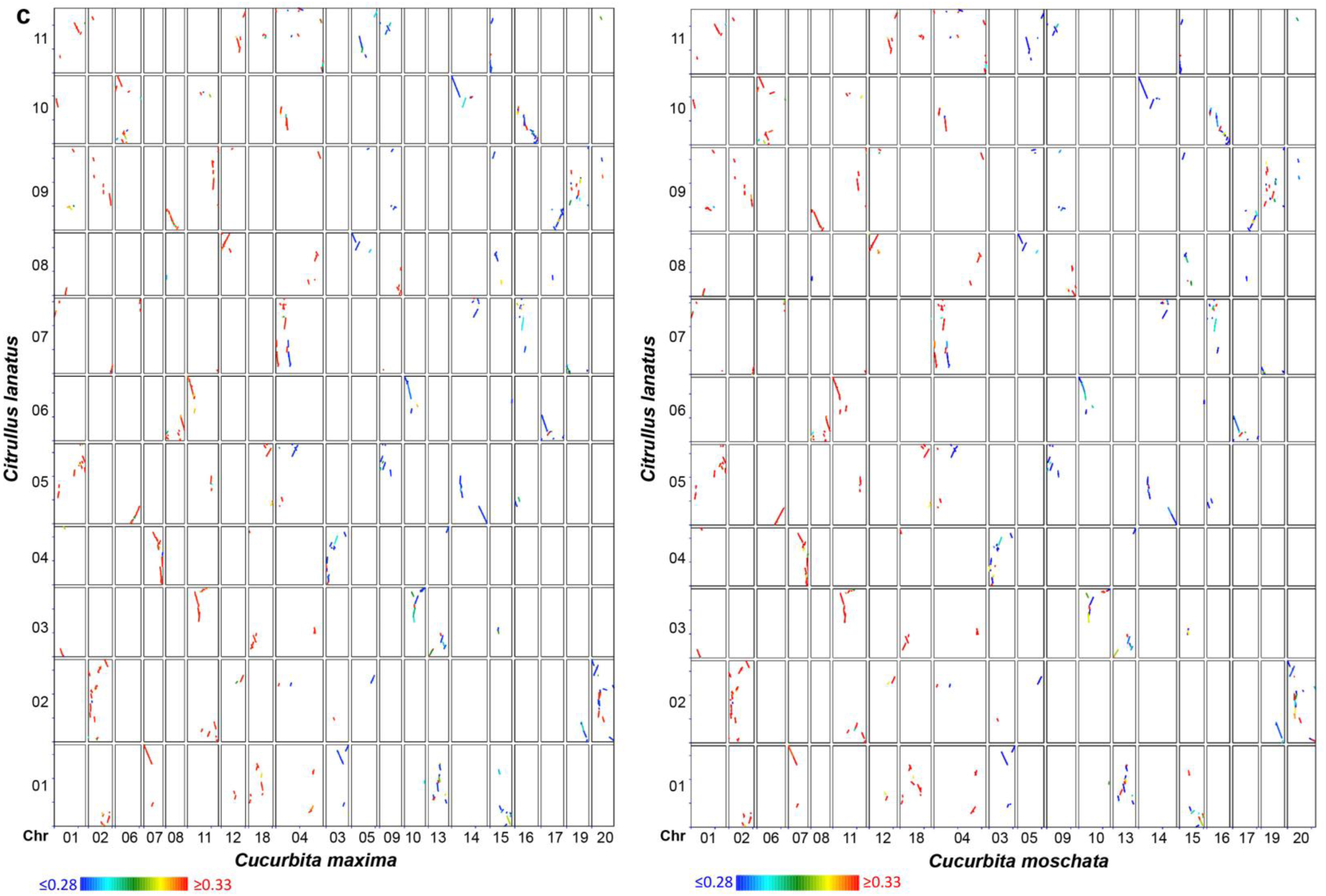
Comparison between genomes of *Cucurbita* and Benincaseae species. (**a**) *C. moschata vs*. melon (*Cucumis melo*). (**b**) *C. maxima* (or *C. moschata*) *vs*. cucumber (*Cucumis sativus*). (**c**) *C. maxima* (or *C. moschata*) *vs*. watermelon (*Citrullus lanatus*). Different *K*s values are marked by different colors.

**Figure S7.**
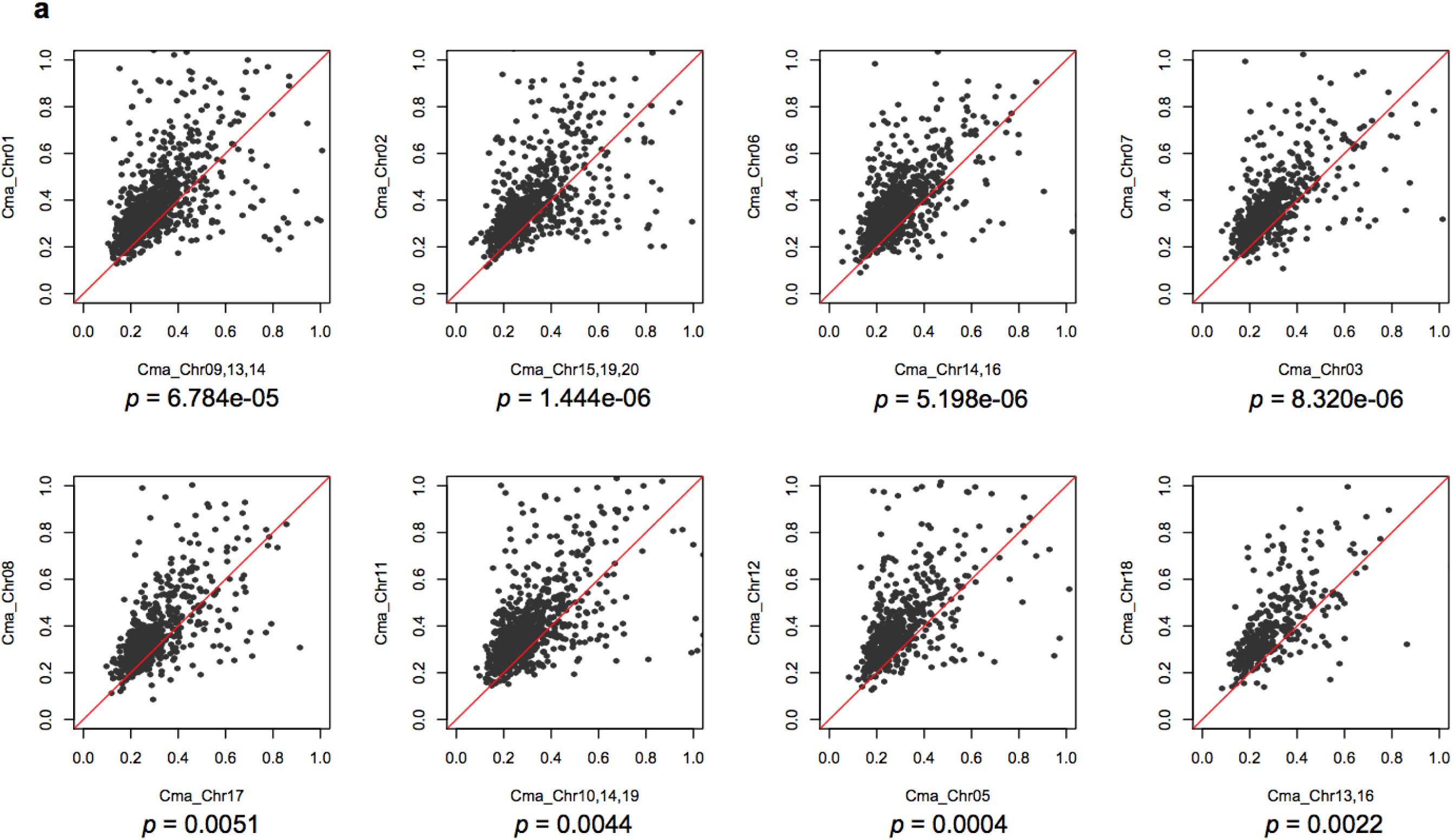
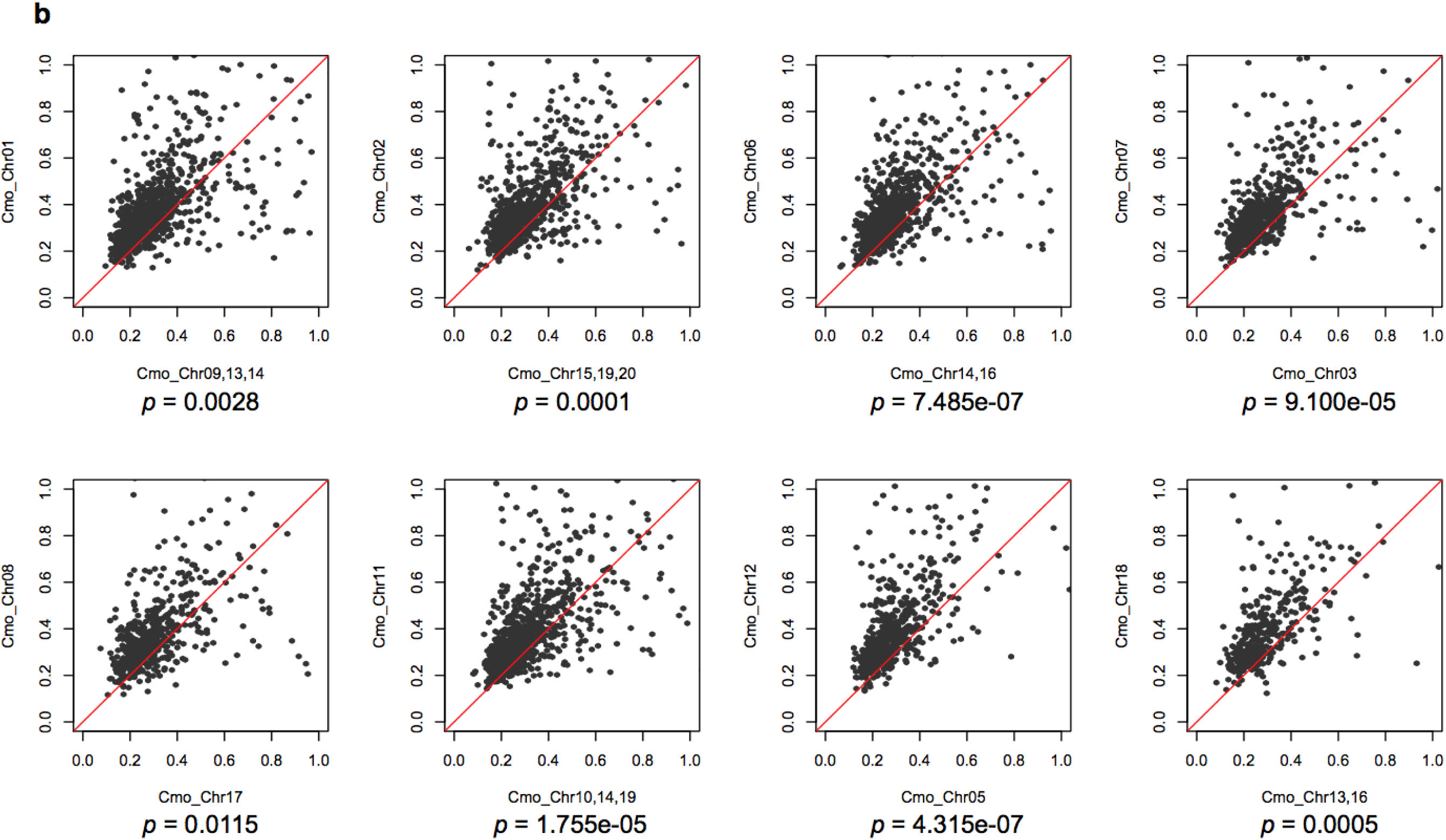
Scatter plots of *Cucurbita*-watermelon *K*s values between pairs of syntenic paralogous genes in *C. maxima* (a) and *C. moschata* (b). *K*s values of the genes in subgenome A (*y* axis) tended to be significantly higher than those of genes in subgenome B (*x* axis). *P-*values of *t*-tests are shown.

**Figure S8.**
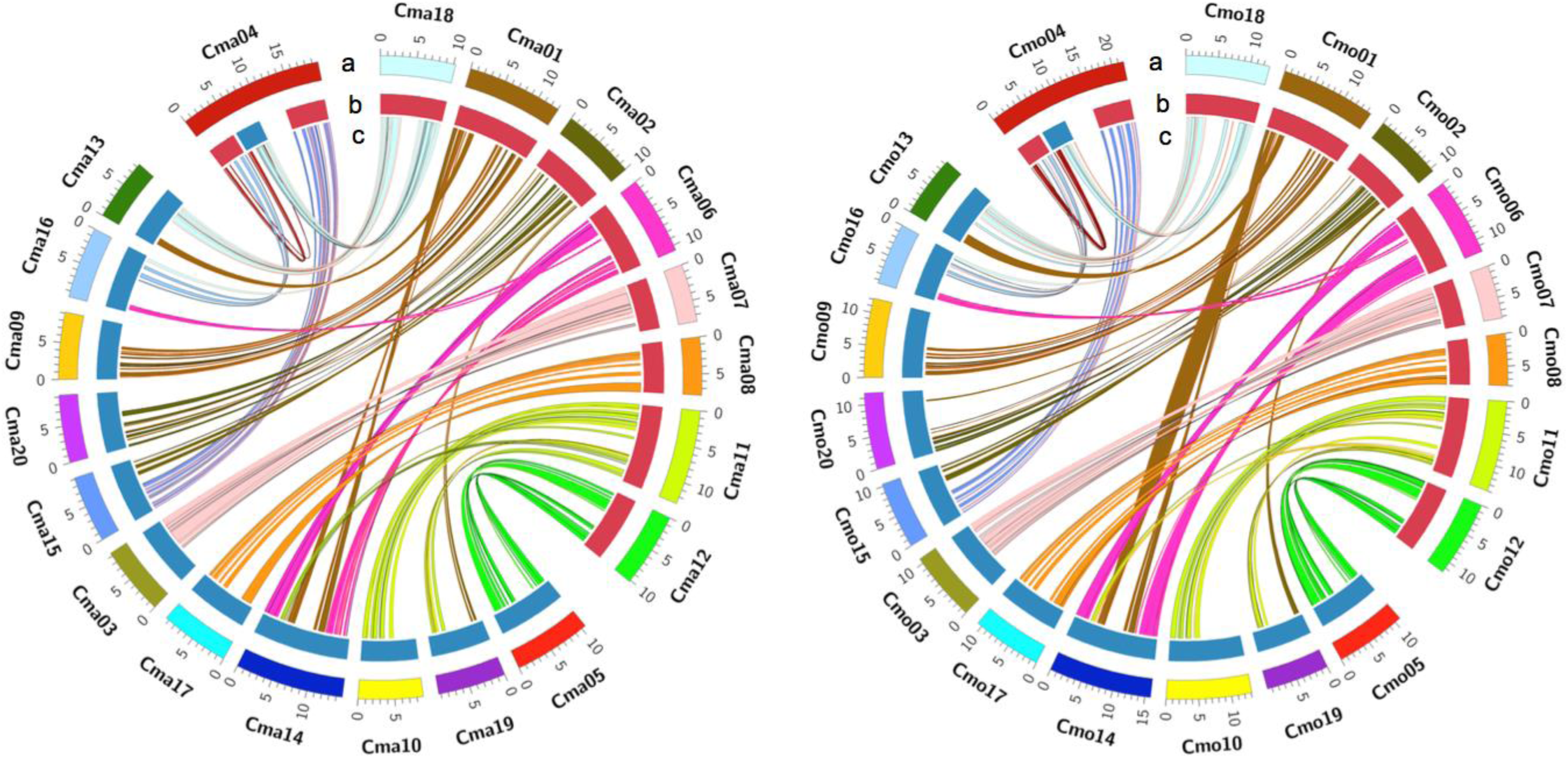
Two subgenomes in *C. maxima* (left) and *C. moschata* (right). (**a**) Ideogram of the 20 pseudochromosomes (in Mb scale). (**b**) Assignment of chromosomes to subgenome A (red) and B (blue). (**c**) Homoeologous regions depicted by connected lines. Each homoeologous block in *Cucurbita* was required to be syntenic to the same region in watermelon. Cma, *C. maxima*. Cmo, *C. moschata*.

**Figure S9.**
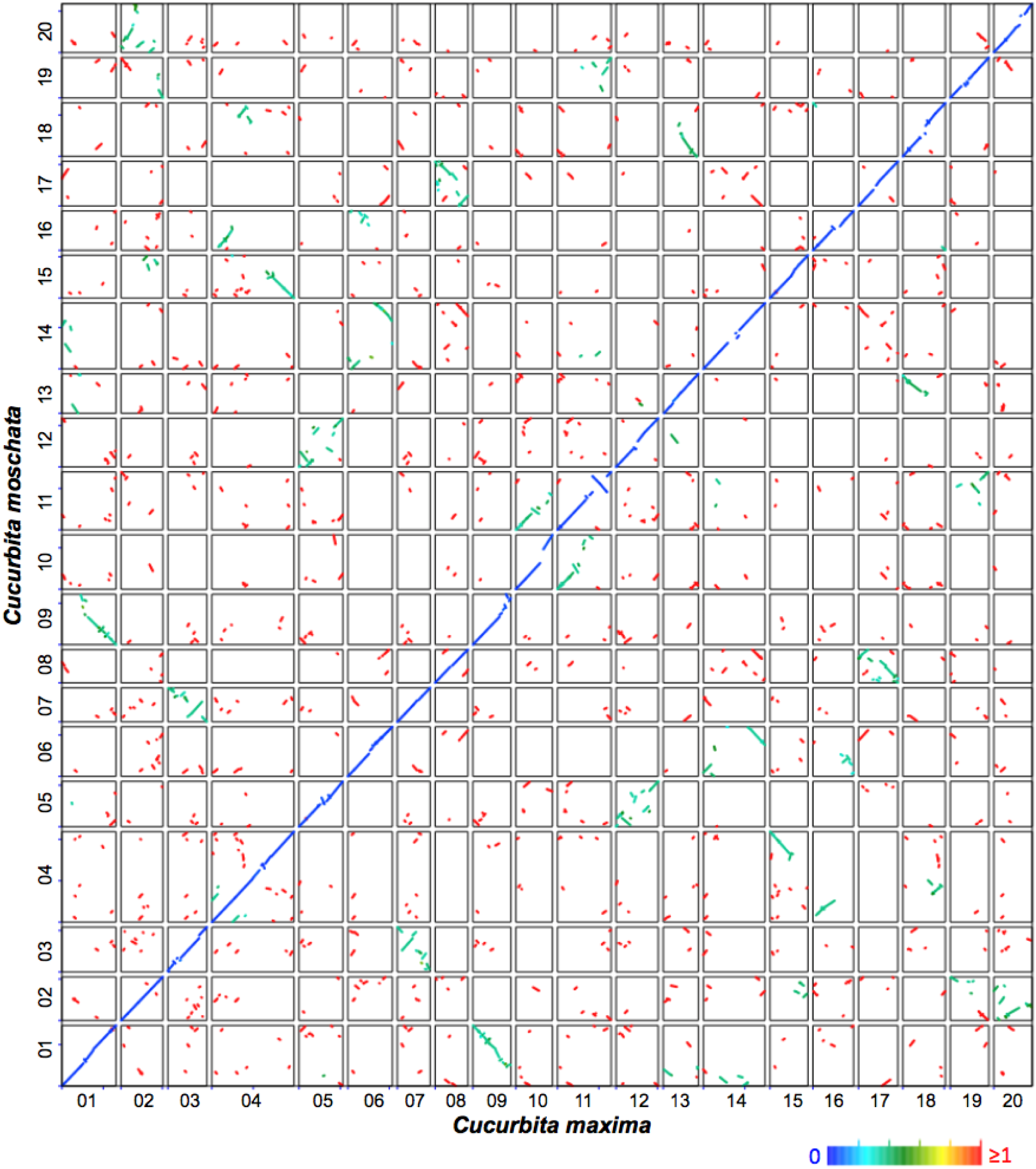
Syntenic dotplots showing the comparison between the genomes of *C. maxima* and *C. moschata*. Orthologous gene pairs are marked in color based on median *K*s values of syntenic blocks.

**Figure S10.**
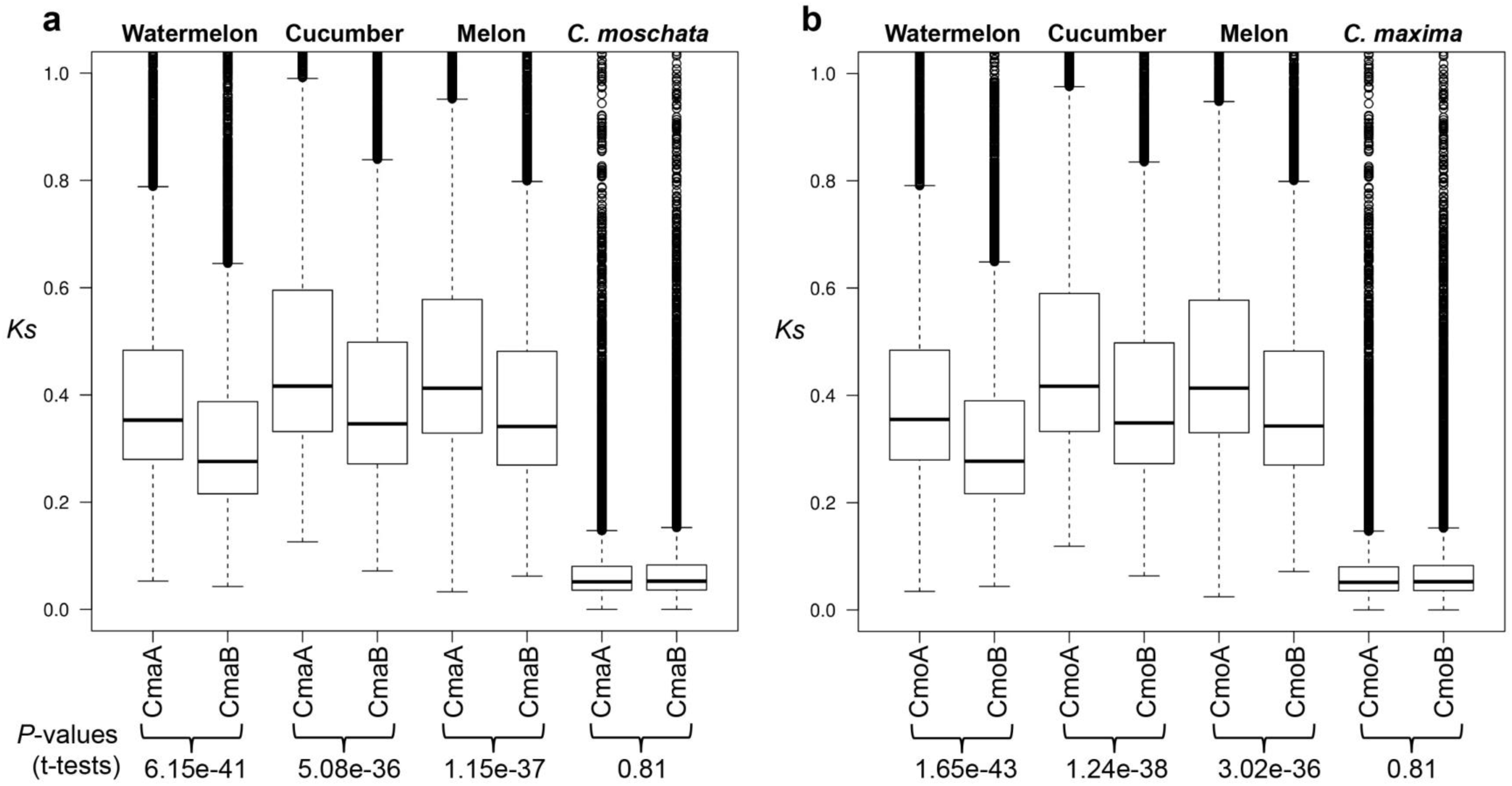
Comparisons of the distribution of *K*s values between the two subgenomes of *C. maxima* (a) and *C. moschata* (b). Cma, *C. maxima*. Cmo, *C. moschata*.

**Figure S11.**
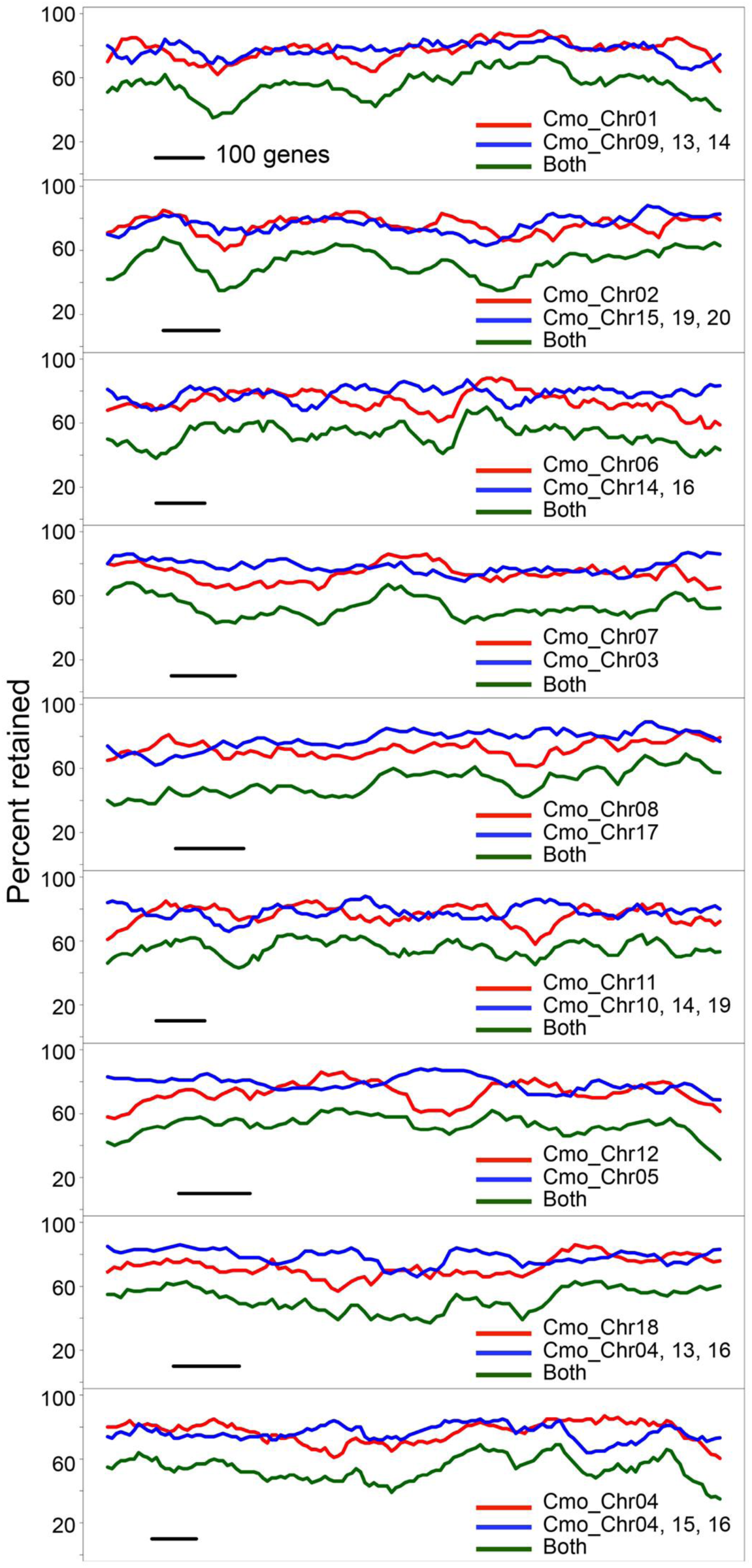
Fractionation in pairs of homoeologous regions in *C. moschata*. Average gene retention percentages are shown in a 100-gene window along each subgenome. Retention of genes in subgenomes A, B and both are shown in red, blue and green, respectively.

**Figure S12.**
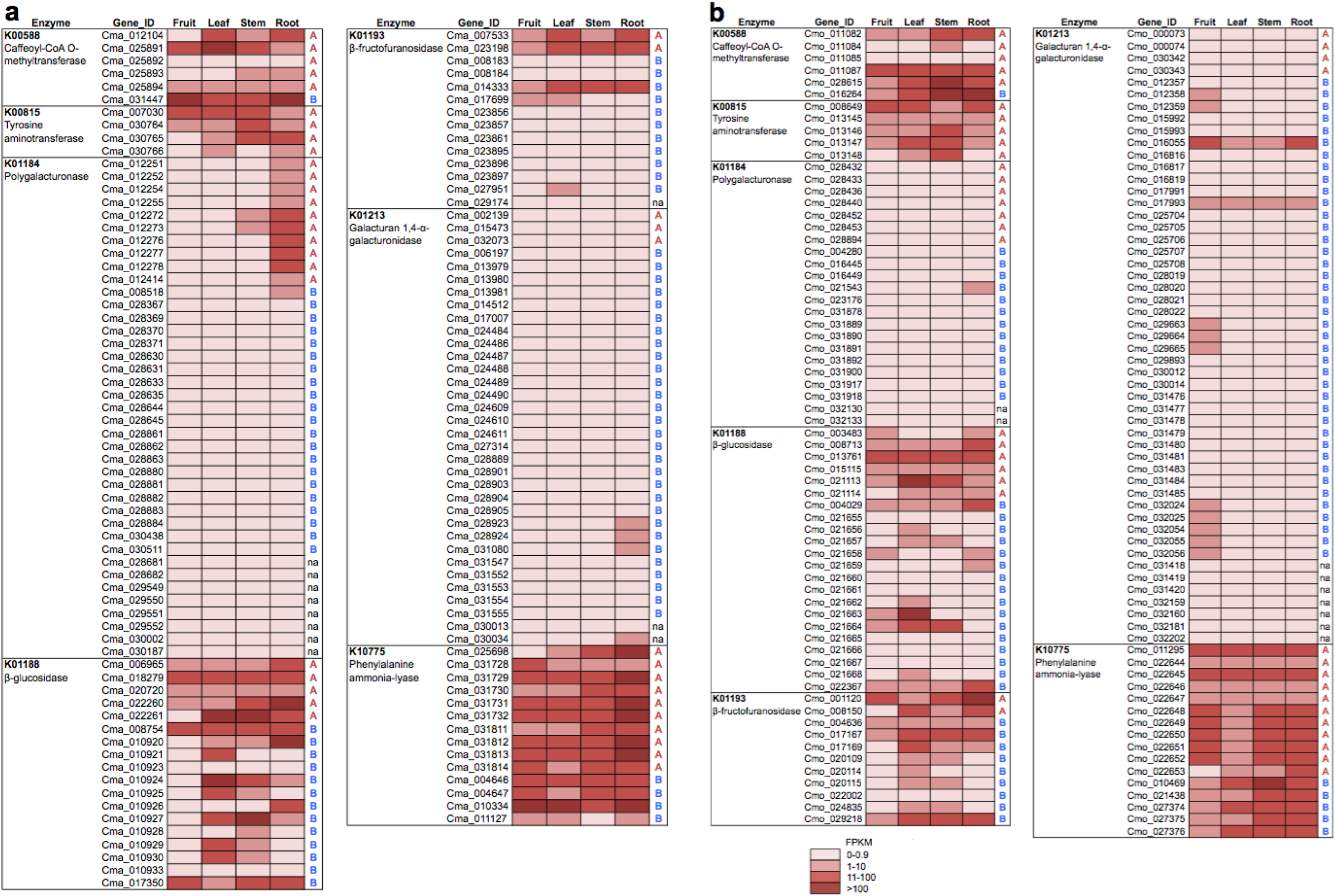
Enzymes encoded by different numbers of genes from subgenomes A and B of *C. maxima* (a) and *C. moschata* (b).

**Figure S13.**
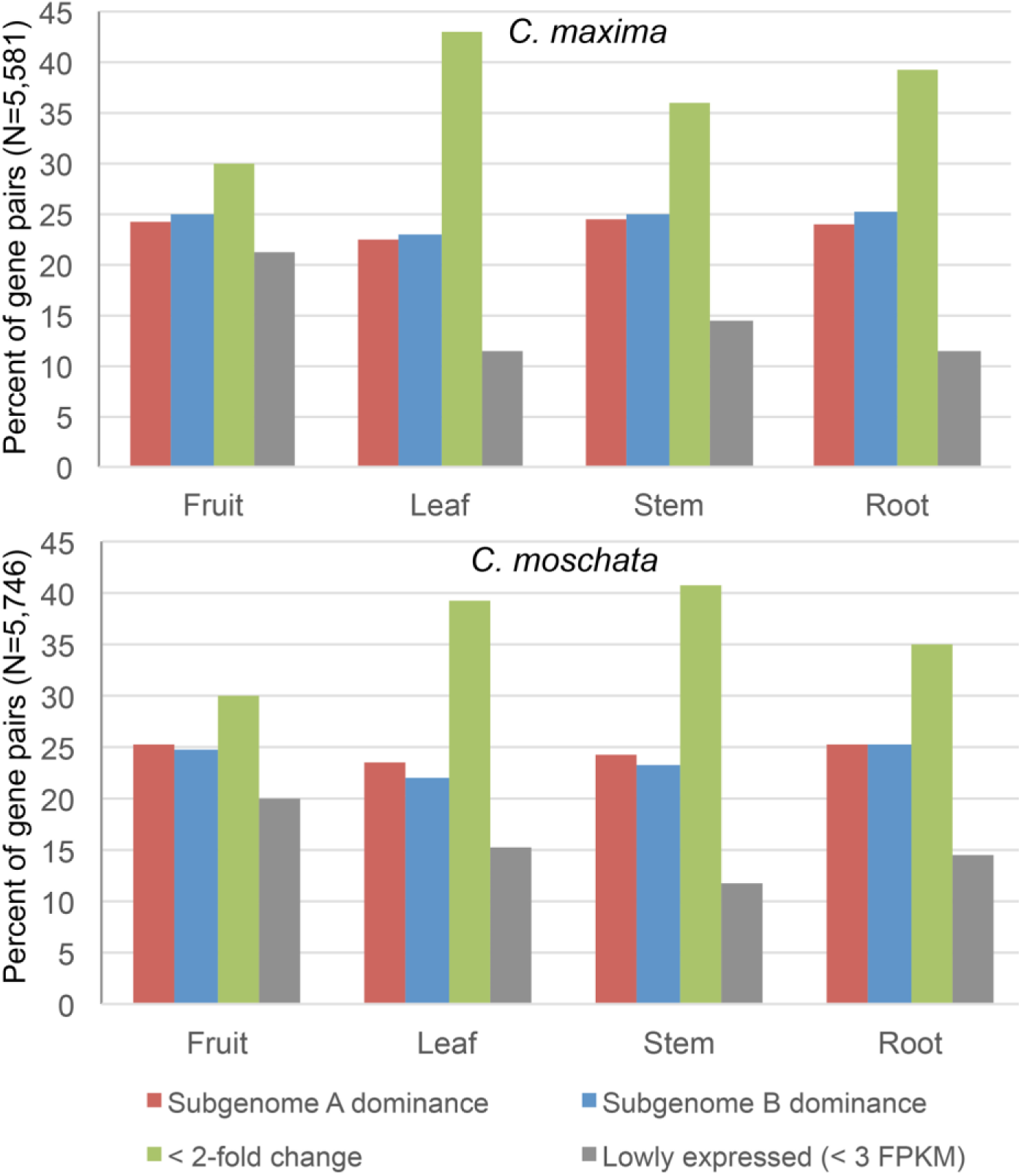
Expression classification of high-confidence homoeologous genes pairs in the two subgenomes of *C. maxima* or *C. moschata* in fruit, leaf, stem and root. Paralogs were considered differentially expressed if one was expressed at least twice that of the other. Lowly expressed genes (< 3 FPKM across all samples) were not used in the comparison.

**Figure S14.**
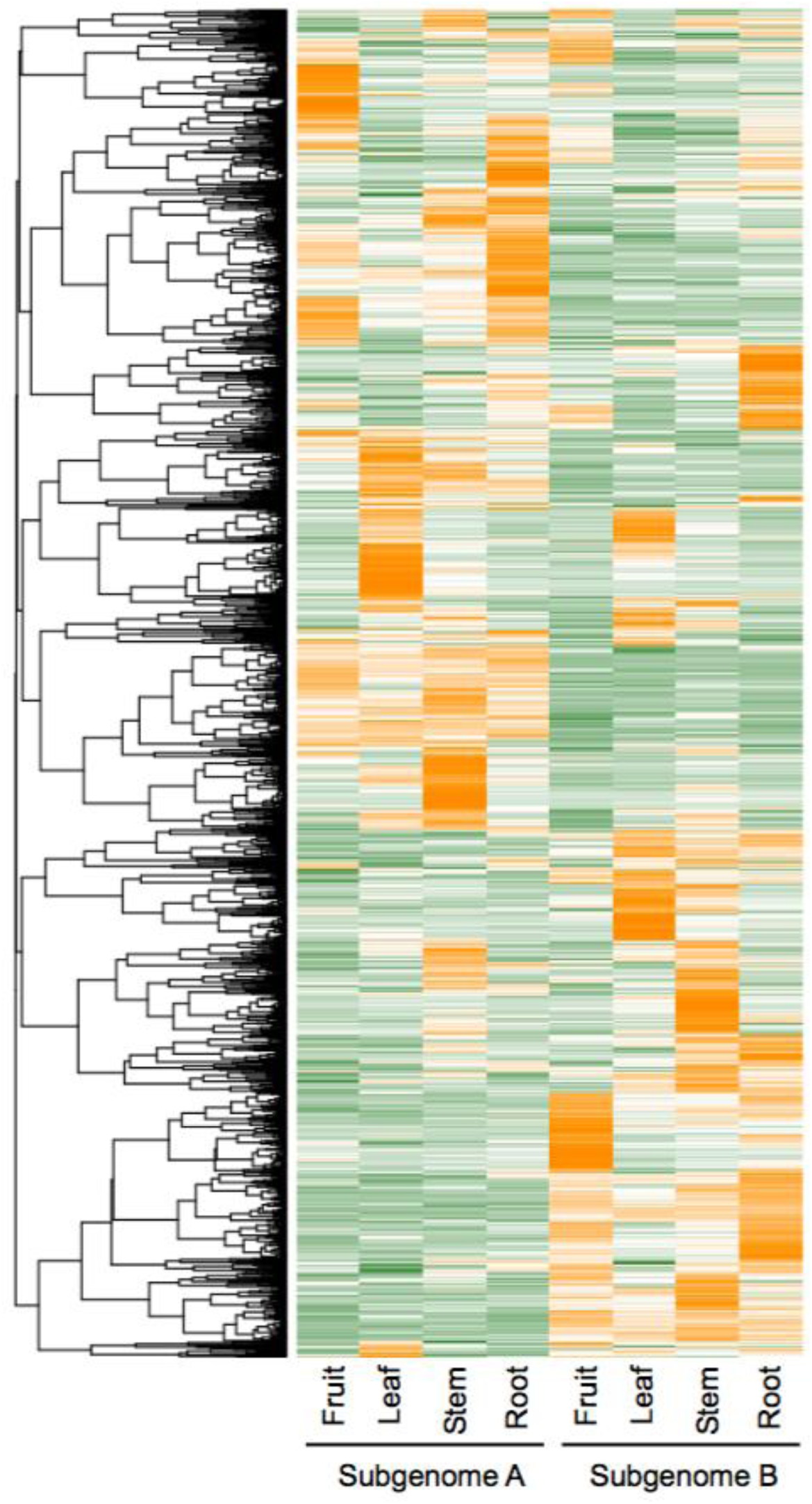
Expression profiles of high-confidence homoeologous gene pairs in the two subgenomes of *C. moschata*. FPKM values of each pair were standardized to have a mean of zero and standard deviation of one across all the samples for the clustering analysis.

**Figure S15.**
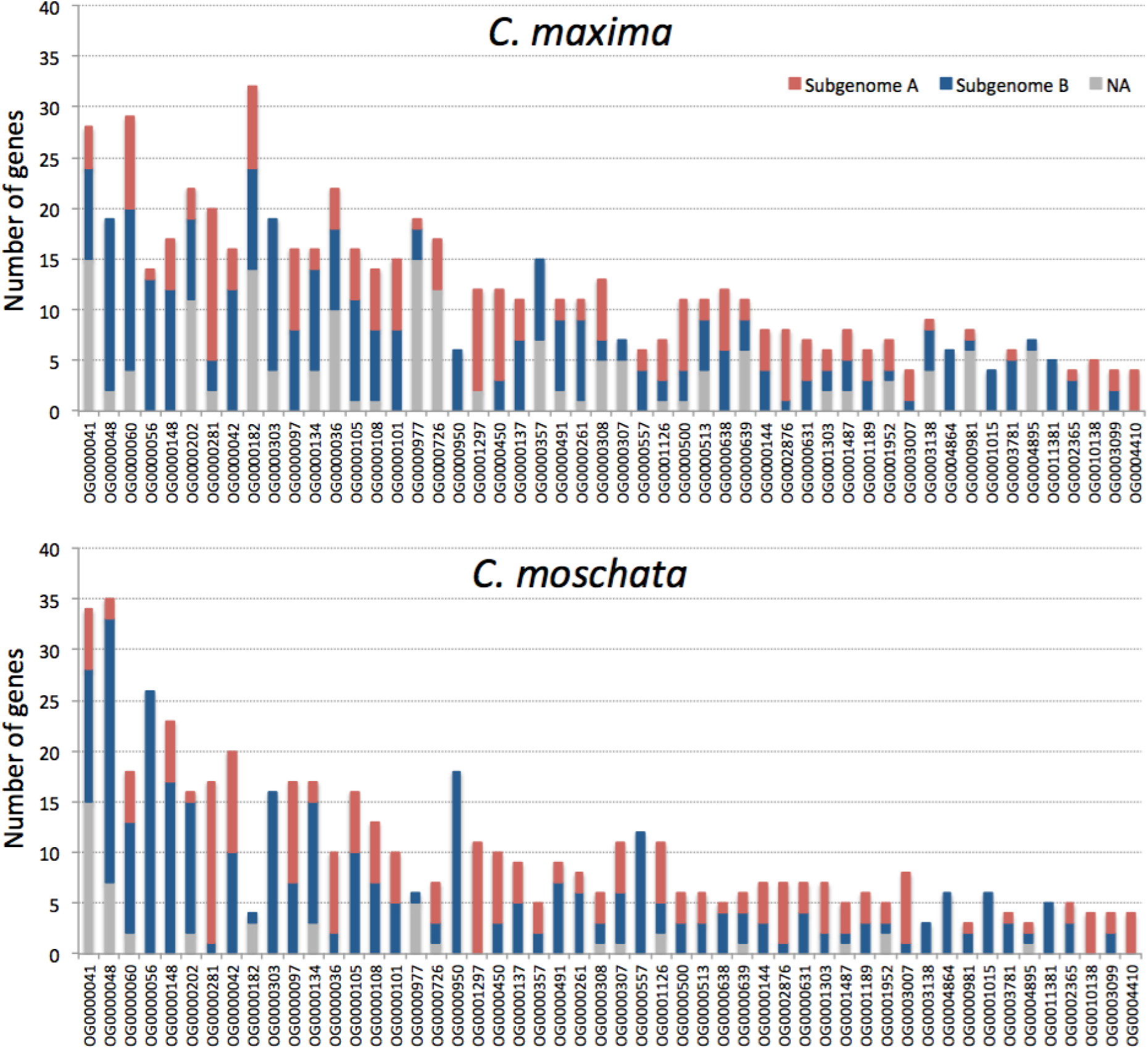
Number of genes in the expanded families in the *Cucurbita* lineage.

**Figure S16.**
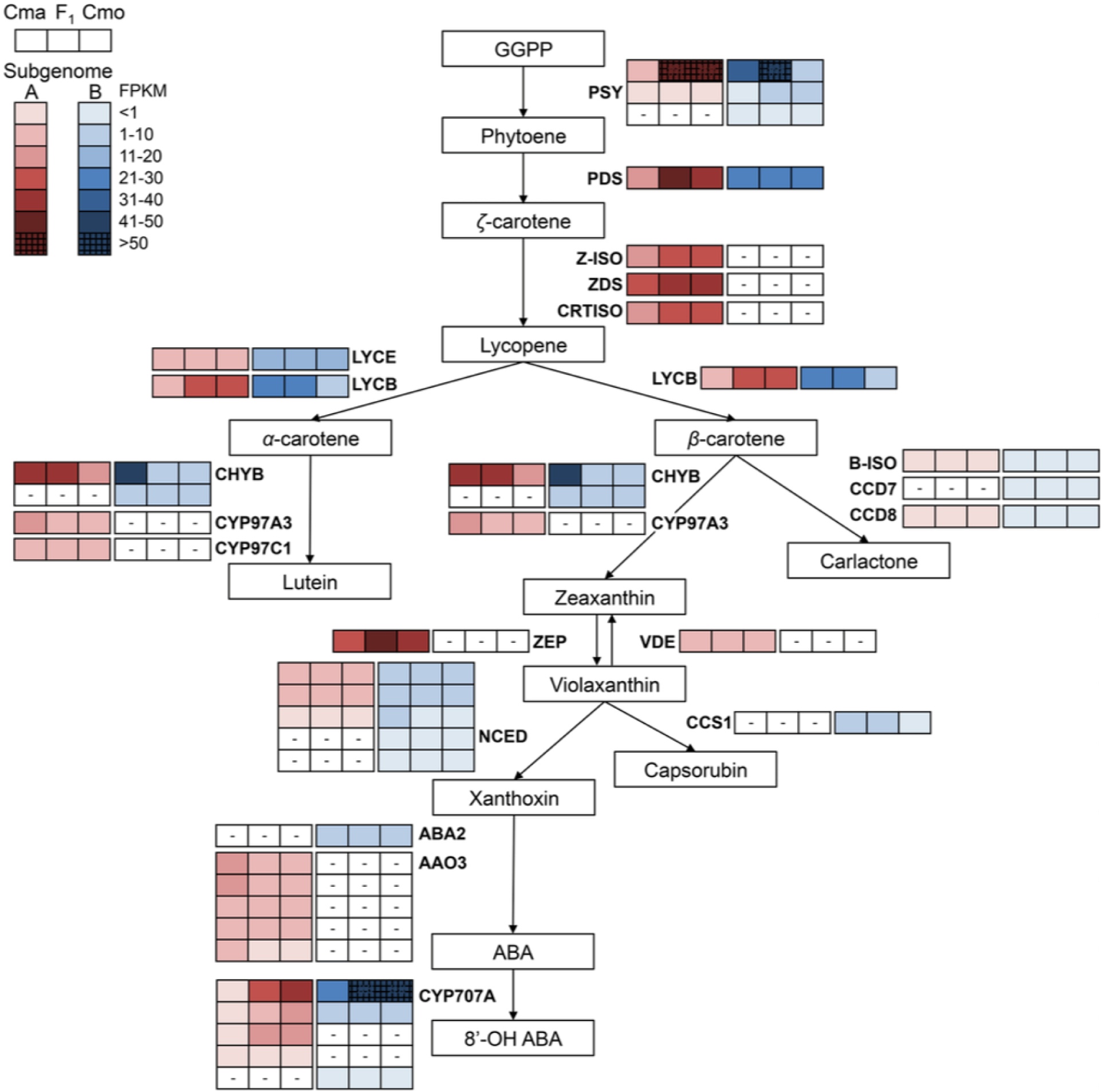
Expression of genes in the carotenoid biosynthetic pathway. The expression levels (FPKM) of homoeologous genes in the fruits of *C. maxima* (Cma), *C. moschata* (Cmo) and the F_1_ hybrid are shown in the same row, with those from subgenome A and B on the left (red) and right (blue), respectively. White boxes with “-” indicate lost homoeologous genes.

## References

1. Bisognin DA. Origin and evolution of cultivated cucurbits. Ciênc. Rural. 2002;32:715–23.

2. Ferriol M, Pico B. Pumpkin and Winter Squash. Handb. Plant Breed. New York, NY: Springer New York; 2008. p. 317–49.

3. Loy JB. Morpho-physiological aspects of productivity and auality in squash and pumpkins (*Cucurbita* spp.). Crit. Rev. Plant Sci. 2004;23:337–63.

4. Nee M. The Domestication of *Cucurbita*. Econ. Bot. 1990;44:56–68.

5. Lee J-M, Kubota C, Tsao SJ, Bie Z, Echevarria PH, Morra L, et al. Current status of vegetable grafting: Diffusion, grafting techniques, automation. Sci. Hortic. 2010;127:93–105.

6. Davis AR, Perkins-Veazie P, Sakata Y, Lopez-Galarza S, Maroto JV, Lee S-G, et al. Cucurbit grafting. Crit. Rev. Plant Sci. 2008;27:50–74.

7. Whitaker T, R R. Squash breeding. In: Baset M, editor. Breed. Veg. Crop. Westport, Connecticut: AVI Publishing Company, Inc.; 1986. p. 209–42.

8. Weeden N. Isozyme studies indicate that the genus *Cucurbita* is an ancient tetraploid. Cucurbit Genet. Coop. Rep. 1984;7:84–5.

9. Singh AK. Cytogenetics and evolution in the Cucurbitaceae. In: Bates D, Robinson R, Jeffrey C, editors. Biol. Util. Cucurbitaceae. Ithaca, New York: Cornell University Press; 1990. p. 10–28.

10. Esteras C, Gomez P, Monforte AJ, Blanca J, Vicente-Dolera N, Roig C, et al. High-throughput SNP genotyping in *Cucurbita pepo* for map construction and quantitative trait loci mapping. BMC Genomics. 2012;13:80.

11. Zhang G, Ren Y, Sun H, Guo S, Zhang F, Zhang J, et al. A high-density genetic map for anchoring genome sequences and identifying QTLs associated with dwarf vine in pumpkin (*Cucurbita maxima* Duch.). BMC Genomics. 2015;16:1101.

12. Simão FA, Waterhouse RM, Ioannidis P, Kriventseva E V., Zdobnov EM. BUSCO: assessing genome assembly and annotation completeness with single-copy orthologs. Bioinformatics. 2015;31:3210–2.

13. Huang S, Li R, Zhang Z, Li L, Gu X, Fan W, et al. The genome of the cucumber, *Cucumis sativus* L. Nat. Genet. 2009;41:1275–81.

14. Garcia-Mas J, Benjak A, Sanseverino W, Bourgeois M, Mir G, González VM, et al. The genome of melon (*Cucumis melo* L.). Proc. Natl. Acad. Sci. U. S. A. 2012;109:11872–7.

15. Guo S, Zhang J, Sun H, Salse J, Lucas WJ, Zhang H, et al. The draft genome of watermelon (*Citrullus lanatus*) and resequencing of 20 diverse accessions. Nat. Genet. 2013;45:51–8.

16. Li Z, Zhang Z, Yan P, Huang S, Fei Z, Lin K. RNA-Seq improves annotation of protein-coding genes in the cucumber genome. BMC Genomics. 2011;12:540.

17. Lin X, Zhang Y, Kuang H, Chen J. Frequent loss of lineages and deficient duplications accounted for low copy number of disease resistance genes in Cucurbitaceae. BMC Genomics. 2013;14:1.

18. Tang H, Woodhouse MR, Cheng F, Schnable JC, Pedersen BS, Conant G, et al. Altered patterns of fractionation and exon deletions in *Brassica rapa* support a two-step model of paleohexaploidy. Genetics. 2012;190:1563–74.

19. Conant GC, Birchler JA, Pires JC. Dosage, duplication, and diploidization: clarifying the interplay of multiple models for duplicate gene evolution over time. Curr. Opin. Plant Biol. 2014;19:91–8.

20. Thomas BC, Pedersen B, Freeling M. Following tetraploidy in an Arabidopsis ancestor, genes were removed preferentially from one homeolog leaving clusters enriched in dose-sensitive genes. Genome Res. 2006;16:934–46.

21. Schnable JC, Springer NM, Freeling M. Differentiation of the maize subgenomes by genome dominance and both ancient and ongoing gene loss. Proc. Natl. Acad. Sci. U. S. A. 2011;108:4069–74.

22. Zhang T, Hu Y, Jiang W, Fang L, Guan X, Chen J, et al. Sequencing of allotetraploid cotton (*Gossypium hirsutum* L. acc. TM-1) provides a resource for fiber improvement. Nat. Biotechnol. 2015;33:531–7.

23. Schnable JC, Wang X, Pires JC, Freeling M. Escape from preferential retention following repeated whole genome duplications in plants. Front. Plant Sci. 2012;3:94.

24. Freeling M, Thomas BC. Gene-balanced duplications, like tetraploidy, provide predictable drive to increase morphological complexity. Genome Res. 2006;16:805–14.

25. Maere S, De Bodt S, Raes J, Casneuf T, Van Montagu M, Kuiper M, et al. Modeling gene and genome duplications in eukaryotes. Proc. Natl. Acad. Sci. U. S. A. 2005;102:5454–9.

26. De Smet R, Van de Peer Y. Redundancy and rewiring of genetic networks following genome-wide duplication events. Curr. Opin. Plant Biol. 2012;15:168–76.

27. Buggs RJA, Wendel JF, Doyle JJ, Soltis DE, Soltis PS, Coate JE. The legacy of diploid progenitors in allopolyploid gene expression patterns. Philos. Trans. R. Soc. B Biol. Sci. 2014;369:20130354–20130354.

28. Flagel L, Udall J, Nettleton D, Wendel J. Duplicate gene expression in allopolyploid *Gossypium* reveals two temporally distinct phases of expression evolution. BMC Biol. 2008;6:16.

29. Yoo M-J, Szadkowski E, Wendel JF. Homoeolog expression bias and expression level dominance in allopolyploid cotton. Heredity. 2013;110:171–80.

30. Liu D, Sun W, Yuan Y, Zhang N, Hayward A, Liu Y, et al. Phylogenetic analyses provide the first insights into the evolution of OVATE family proteins in land plants. Ann. Bot. 2014;113:1219–33.

31. Ha M, Kim E-D, Chen ZJ. Duplicate genes increase expression diversity in closely related species and allopolyploids. Proc. Natl. Acad. Sci. U. S. A. 2009;106:2295–300.

32. Wittkopp PJ, Haerum BK, Clark AG. Regulatory changes underlying expression differences within and between *Drosophila* species. Nat. Genet. 2008;40:346–50.

33. Lemos B, Araripe LO, Fontanillas P, Hartl DL. Dominance and the evolutionary accumulation of *cis*- and *trans*-effects on gene expression. Proc. Natl. Acad. Sci. U. S. A. 2008;105:14471–6.

34. Bell GDM, Kane NC, Rieseberg LH, Adams KL. RNA-Seq analysis of allele-specific expression, hybrid effects, and regulatory divergence in hybrids compared with their parents from natural populations. Genome Biol. Evol. 2013;5:1309–23.

35. Nakkanong K, Yang JH, Zhang MF. Carotenoid accumulation and carotenogenic gene expression during fruit development in novel interspecific inbred squash lines and their parents. J. Agric. Food Chem. 2012;60:5936–44.

36. Pearson O, Hopp R, Bohn G. Notes on species crosses in *Cucurbita*. Proc. Am. Soc. Hortic. Sci. 1951;57:310–322.

37. Ara N, Yang JH, Hu ZY, Zhang MF. Determining heat tolerance of interspecific (*Cucurbita maxima* × *Cucurbita moschata*) inbred line of squash “Maxchata” and its parents through photosynthetic response. Tarım Bilim. Derg.-J. Agric. Sci. 2013;19.

38. Chen ZJ. Genomic and epigenetic insights into the molecular bases of heterosis. Nat. Rev. Genet. 2013;14:471–82.

39. Groszmann M, Gonzalez-Bayon R, Lyons RL, Greaves IK, Kazan K, Peacock WJ, et al. Hormone-regulated defense and stress response networks contribute to heterosis in *Arabidopsis* F_1_ hybrids. Proc. Natl. Acad. Sci. 2015;112:E6397–406.

40. Fujimoto R, Taylor JM, Shirasawa S, Peacock WJ, Dennis ES. Heterosis of *Arabidopsis* hybrids between C24 and Col is associated with increased photosynthesis capacity. Proc. Natl. Acad. Sci. U. S. A. 2012;109:7109–14.

41. Renner SS, Schaefer H. Phylogeny and evolution of the Cucurbitaceae. In: Grumet R, Katzir N, Garcia-Mas J, editors. Genet. genomics Cucurbitaceae. New York, NY: Springer New York; 2016. p. 1–11.

42. Li F, Fan G, Lu C, Xiao G, Zou C, Kohel RJ, et al. Genome sequence of cultivated Upland cotton (*Gossypium hirsutum* TM-1) provides insights into genome evolution. Nat. Biotechnol. 2015;33:524–30.

43. Renny-Byfield S, Wendel JF. Doubling down on genomes: polyploidy and crop plants. Am. J. Bot. 2014;101:1711–25.

44. Session AM, Uno Y, Kwon T, Chapman JA, Toyoda A, Takahashi S, et al. Genome evolution in the allotetraploid frog *Xenopus laevis*. Nature. 2016;538:336–43.

45. Zhang H, Bian Y, Gou X, Dong Y, Rustgi S, Zhang B, et al. Intrinsic karyotype stability and gene copy number variations may have laid the foundation for tetraploid wheat formation. Proc. Natl. Acad. Sci. U. S. A. 2013;110:19466–71.

46. Panchy N, Lehti-Shiu M, Shiu S-H. Evolution of Gene Duplication in Plants. Plant Physiol. 2016;171:2294–316.

47. Garsmeur O, Schnable JC, Almeida A, Jourda C, D’Hont A, Freeling M. Two evolutionarily distinct classes of paleopolyploidy. Mol. Biol. Evol. 2014;31:448–54.

48. Gill N, Findley S, Walling JG, Hans C, Ma J, Doyle J, et al. Molecular and chromosomal evidence for allopolyploidy in soybean. Plant Physiol. 2009;151:1167–74.

49. Azevedo-Meleiro CH, Rodriguez-Amaya DB. Qualitative and quantitative differences in carotenoid composition among *Cucurbita moschata*, *Cucurbita maxima*, and *Cucurbita pepo*. J. Agric. Food Chem. 2007;55:4027–33.

50. Zhong S, Joung J-G, Zheng Y, Chen Y -r., Liu B, Shao Y, et al. High-throughput Illumina strand-specific RNA sequencing library preparation. Cold Spring Harb. Protoc. 2011;2011:pdb.prot5652-prot5652.

51. Bolger AM, Lohse M, Usadel B. Trimmomatic: a flexible trimmer for Illumina sequence data. Bioinformatics. 2014;30:2114–20.

52. Morgan M, Anders S, Lawrence M, Aboyoun P, Pages H, Gentleman R. ShortRead: a bioconductor package for input, quality assessment and exploration of high-throughput sequence data. Bioinformatics. 2009;25:2607–8.

53. Marcais G, Yorke JA, Zimin A. QuorUM: An error corrector for Illumina reads. PLoS One. 2015;10:e0130821.

54. Luo R, Liu B, Xie Y, Li Z, Huang W, Yuan J, et al. SOAPdenovo2: an empirically improved memory-efficient short-read de novo assembler. Gigascience. 2012;1:18.

55. Walker BJ, Abeel T, Shea T, Priest M, Abouelliel A, Sakthikumar S, et al. Pilon: an integrated tool for comprehensive microbial variant detection and genome assembly improvement. Wang J, editor. PLoS One. 2014;9:e112963.

56. Elshire RJ, Glaubitz JC, Sun Q, Poland JA, Kawamoto K, Buckler ES, et al. A robust, simple genotyping-by-sequencing (GBS) approach for high diversity species. Orban L, editor. PLoS One. 2011;6:e19379.

57. Glaubitz JC, Casstevens TM, Lu F, Harriman J, Elshire RJ, Sun Q, et al. TASSEL-GBS: A high capacity genotyping by sequencing analysis pipeline. Tinker NA, editor. PLoS One. 2014;9:e90346.

58. Li H, Durbin R. Fast and accurate short read alignment with Burrows-Wheeler transform. Bioinformatics. 2009;25:1754–60.

59. Wu Y, Bhat PR, Close TJ, Lonardi S. Efficient and accurate construction of genetic linkage maps from the minimum spanning tree of a graph. PLoS Genet. 2008;4:e1000212.

60. Ellinghaus D, Kurtz S, Willhoeft U. LTRharvest, an efficient and flexible software for de novo detection of LTR retrotransposons. BMC Bioinformatics. 2008;9:18.

61. Han Y, Wessler SR. MITE-Hunter: a program for discovering miniature inverted-repeat transposable elements from genomic sequences. Nucleic Acids Res. 2010;38:e199–e199.

62. UniProt Consortium. Ongoing and future developments at the Universal Protein Resource. Nucleic Acids Res. 2011;39:D214–9.

63. Feschotte C, Keswani U, Ranganathan N, Guibotsy ML, Levine D. Exploring repetitive DNA landscapes using REPCLASS, a tool that automates the classification of transposable elements in eukaryotic genomes. Genome Biol. Evol. 2009;1:205–20.

64. Cantarel BL, Korf I, Robb SMC, Parra G, Ross E, Moore B, et al. MAKER: An easy-to-use annotation pipeline designed for emerging model organism genomes. Genome Res. 2007;18:188–96.

65. Korf I. Gene finding in novel genomes. BMC Bioinformatics. 2004;5:59.

66. Stanke M, Tzvetkova A, Morgenstern B. AUGUSTUS at EGASP: using EST, protein and genomic alignments for improved gene prediction in the human genome. Genome Biol. 2006;7:S 11.

67. Grabherr MG, Haas BJ, Yassour M, Levin JZ, Thompson DA, Amit I, et al. Full-length transcriptome assembly from RNA-Seq data without a reference genome. Nat. Biotechnol. 2011;29:644–52.

68. Haas BJ, Delcher AL, Mount SM, Wortman JR, Smith RK, Hannick LI, et al. Improving the Arabidopsis genome annotation using maximal transcript alignment assemblies. Nucleic Acids Res. 2003;31:5654–66.

69. The Arabidopsis Genome Initiative. Analysis of the genome sequence of the flowering plant *Arabidopsis thaliana*. Nature. 2000;408:796–815.

70. Iwata H, Gotoh O. Benchmarking spliced alignment programs including Spaln2, an extended version of Spaln that incorporates additional species-specific features. Nucleic Acids Res. 2012;40:e161.

71. Urasaki N, Takagi H, Natsume S, Uemura A, Taniai N, Miyagi N, et al. Draft genome sequence of bitter gourd (*Momordica charantia*), a vegetable and medicinal plant in tropical and subtropical regions. DNA Res. 2016;24:51-58.

72. Jones P, Binns D, Chang H-Y, Fraser M, Li W, McAnulla C, et al. InterProScan 5: genome-scale protein function classification. Bioinformatics. 2014;30:1236–40.

73. Conesa A, Gotz S, Garcia-Gomez JM, Terol J, Talon M, Robles M. Blast2GO: a universal tool for annotation, visualization and analysis in functional genomics research. Bioinformatics. 2005;21:3674–6.

74. Zheng Y, Jiao C, Sun H, Rosli HG, Pombo MA, Zhang P, et al. iTAK: a program for genome-wide prediction and classification of plant transcription factors, transcriptional regulators, and protein kinases. Mol. Plant. 2016;9:1667–70.

75. Han M V., Thomas GWC, Lugo-Martinez J, Hahn MW. Estimating gene gain and loss rates in the presence of error in genome assembly and annotation using CAFE 3. Mol. Biol. Evol. 2013;30:1987–97.

76. Emms DM, Kelly S. OrthoFinder: solving fundamental biases in whole genome comparisons dramatically improves orthogroup inference accuracy. Genome Biol. 2015;16:157.

77. Zhang J, Nielsen R, Yang Z. Evaluation of an improved branch-site likelihood method for detecting positive selection at the molecular level. Mol. Biol. Evol. 2005;22:2472–9.

78. Yang Z. Likelihood ratio tests for detecting positive selection and application to primate lysozyme evolution. Mol. Biol. Evol. 1998;15:568–73.

79. Wang Y, Tang H, Debarry JD, Tan X, Li J, Wang X, et al. MCScanX: a toolkit for detection and evolutionary analysis of gene synteny and collinearity. Nucleic Acids Res. 2012;40:e49.

80. Yang Z. PAML: a program package for phylogenetic analysis by maximum likelihood. Bioinformatics. 1997;13:555–6.

81. Guindon S, Dufayard JF, Lefort V, Anisimova M, Hordijk W, Gascuel O. New algorithms and methods to estimate maximum-likelihood phylogenies: assessing the performance of PhyML 3.0. Syst. Biol. 2010;59:307–21.

82. Bouckaert R, Heled J, Kühnert D, Vaughan T, Wu C-H, Xie D, et al. BEAST 2: a software platform for Bayesian evolutionary analysis. Prlic A, editor. PLoS Comput. Biol. 2014;10:e1003537.

83. Yang Z. The BPP program for species tree estimation and species delimitation. Curr. Zool. 2015;61:854–65.

84. Wikström N, Savolainen V, Chase MW. Evolution of the angiosperms: calibrating the family tree. Proc. R. Soc. London B Biol. Sci. 2001;268.

85. Kim D, Langmead B, Salzberg SL. HISAT: a fast spliced aligner with low memory requirements. Nat. Methods. 2015;12:357–60.

86. Anders S, Huber W. Differential expression analysis for sequence count data. Genome Biol. 2010;11:R106.

87. Kiełbasa SM, Wan R, Sato K, Horton P, Frith MC. Adaptive seeds tame genomic sequence comparison. Genome Res. 2011;21:487–93.

88. McManus CJ, Coolon JD, Duff MO, Eipper-Mains J, Graveley BR, Wittkopp PJ. Regulatory divergence in *Drosophila* revealed by mRNA-seq. Genome Res. 2010;20:816–25.

